# Discovery of novel potent, cell-permeable inhibitors of *P. falciparum* cGMP-dependent protein kinase

**DOI:** 10.1101/2021.08.24.457553

**Authors:** Rammohan R. Yadav Bheemanaboina, Mariana Lozano Gonzalez, Shams Ul Mahmood, Tyler Eck, Tamara Kreiss, Mariana Laureano de Souza, Samantha O. Aylor, Alison Roth, Patricia Lee, Brandon S. Pybus, Purnima Bhanot, Dennis J. Colussi, Wayne E. Childers, John Gordon, John J. Siekierka, David P. Rotella

## Abstract

The discovery of new targets for treatment of malaria advanced with the demonstration that orally administered inhibitors of *Plasmodium falciparum* cGMP-dependent protein kinase (PfPKG) could clear infection in a murine model. This enthusiasm was tempered by unsatisfactory safety and/or pharmacokinetic issues found with these chemotypes. To address the urgent need for new scaffolds, we recently reported the discovery and optimization of novel, potent isoxazole-based PfPKG inhibitors that lacked any obvious safety warnings. This manuscript presents representative *in vitro* ADME, hERG characterization and cell-based antiparasitic activity of these PfPKG inhibitors. We also report the discovery and structure-activity relationships of a new series with good potency, low hERG activity and cell-based anti-parasitic activity comparable to a literature standard.

---

The emergence of artemisinin-resistant *P. falciparum* in Africa {Uwimana, 2020 #287;Uwimana, 2021 #286} and the slowing decline in deaths from malaria {Organization, 2018 #248} are compelling reminders of the need to identify new targets for malaria prophylaxis and treatment {Burrows, 2017 #221}. *Plasmodium’s* pre-erythrocytic cycle is an attractive point for therapeutic attack because of the very low parasite burden compared to other life cycle stages. Drugs that target pre-erythrocytic stages are an essential component of the anti-malarial effort because a decrease in liver infection by sporozoites significantly reduces severity and incidence of malaria {Alonso, 2005 #101}. There is a dearth of appropriate candidates for this product profile in the global drug development portfolio {Burrows, 2017 #221} the discovery and development of drugs for malaria chemoprotection is relatively neglected compared to other target candidate profiles {Wells, 2015 #195}.

*Plasmodium falciparum* cGMP-dependent protein kinase (PfPKG) is essential in pre-erythrocytic, asexual and sexual stages of the parasite {Hopp, 2012 #169;Alam, 2015 #183;Brochet, 2014 #182}. Baker and co-workers described the discovery and optimization of an orally bioavailable, potent, selective small molecule inhibitor (**1**, figure 1). This imidazopyridine cleared infection at a dose of 10 mg/kg orally in a SCID mouse model.^1-3^

**Figure 1:**
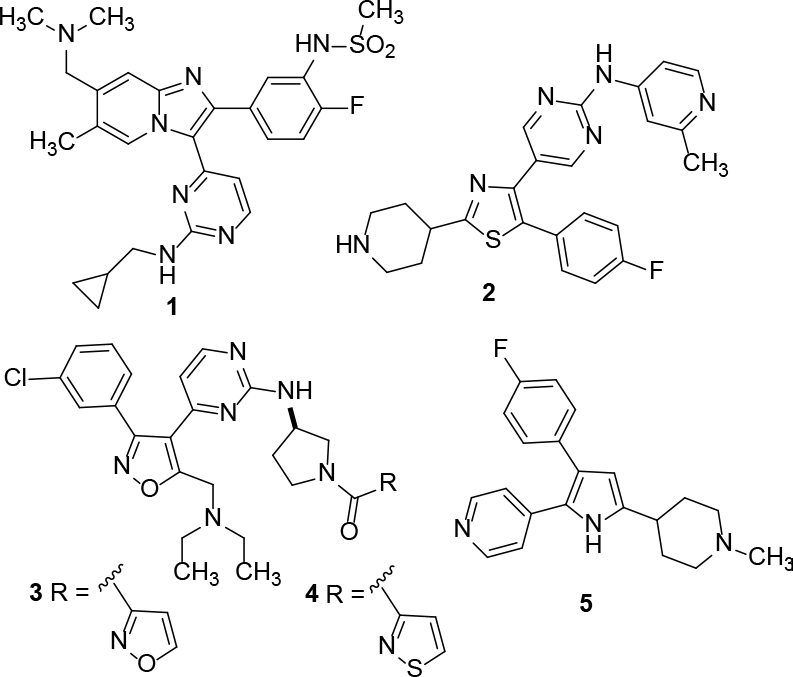
PfPKG Inhibitors

A subsequent report that unspecified examples in this imidazopyridine series were Ames positive, limited progression of the scaffold.^4^ The Baker group disclosed trisubstituted thiazoles such as **2** (figure 1) that exhibited rapid killing of *P. falciparum* in culture.^5^ Interestingly, this desirable property was independent of PfPKG inhibition and proteomic experiments suggested that inhibition of a serine/arginine protein kinase SRPK2 was a key contributor to rapid parasite killing, comparable to artesunate, a recognized standard. Examples in this series of thiazoles, including **2**, showed single digit micromolar hERG activity and/or *in vitro* metabolic instability limiting their use in more advanced studies.

As alternatives to these scaffolds, we previously reported the discovery and initial optimization of a distinct isoxazole chemotype exemplified by **3** and **4** (figure 1).^6^ These compounds showed enzymatic potency comparable to known pyrrole **5**^7^ *in vitro* against PfPKG, and were not active against human PKG or the T618Q mutant PfPKG^8^ at 10 μM. *Plasmodium* mutants with this enzyme retain the ability to proceed through the life cycle and demonstrate lower sensitivity to PfPKG inhibitors such as **5**.^8^ This result stimulated evaluation of the cell-based anti-parasitic activity and *in vitro* ADME properties of these inhibitors.

These results are included in this report, along with the discovery and initial exploration of a second chemotype that displays cellular activity comparable to **5** and the *in vitro* ADME characterization of selected examples in this new template.

Encouraged by the potent enzymatic activity of **3** and **4** (IC_50_s ∼20 nM)^6^, along with excellent selectivity for human PKG and the T618Q mutant, both molecules were examined to cellular studies for anti-parasitic efficacy to reduce infectivity of *P. berghei* sporozoites. It was disappointing to observe that both compounds showed weak or no activity at 2 and 10 μM in this assay where **5** clearly shows greater efficacy (Table 1). These results were accompanied by poor metabolic stability in both murine and human liver microsomes, with half-lives less than 2 minutes for both **3** and **4** (Table 2). Metabolite ID studies suggested that the diethylamino moiety was a likely site of oxidative transformation, leading to a mixture of N-oxidized and N-dealkylated products. We hypothesized that the poor cellular activity of **3** and **4** was associated with their comparatively high molecular weight (> 520 g/mol), in spite of acceptable water solubility (≥ 70 μM/pH 7.4 buffer) that limited their passive diffusion.

**Table 1:**
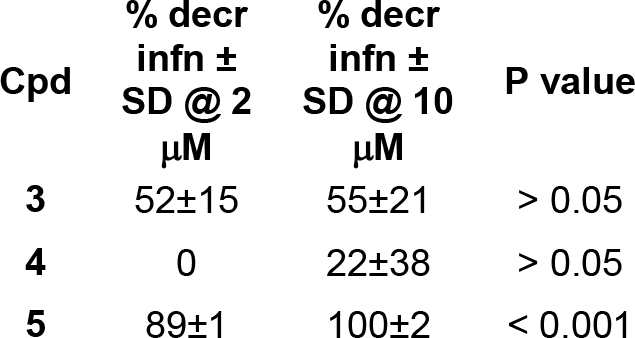
Isoxazole *P. berghei* infectivity assay

**Table 2:** *In vitro* ADME data

| Cpd | HLM<br>t<sub>1/2</sub><br>(min) | MLM<br>t<sub>1/2</sub><br>(min) | H<sub>2</sub>O<br>sol<br>(μM)<br>pH<br>7.4 | 3A4<br>IC<sub>50</sub><br>(μM) | 2C9<br>IC<sub>50</sub><br>(μM) | 2D6<br>IC<sub>50</sub><br>(μM) | hERG<br>%<br>inh.<br>@ 10<br>μM |
| --- | --- | --- | --- | --- | --- | --- | --- |
| **3** | < 2 | < 2 | 83 | 0.41 | 8 | 4.6 | 18 |
| **4** | < 2 | < 2 | 68 | 0.35 | 5.8 | 7.1 | 13 |
| **10a** | < 2 | < 2 | 200 | 0.94 | > 10 | > 10 | NT |
| **14a** | 4.8 | < 2 | 92 | 0.29 | > 10 | > 10 | 1 |
| **14b** | 3.7 | < 2 | 28 | 0.19 | 2.8 | > 10 | 9 |

Efforts to address the dual issues of poor metabolic stability and comparatively high molecular weight with this isoxazole scaffold were unsuccessful. Attention was turned to a second chemotype, imidazole **6** (Figure 2) that was identified in the original focused screen that produced the isoxazole series. We were attracted to the comparatively low molecular weight of this chemotype and the potential for optimization at multiple positions that could result in improved properties. Additionally, the synthesis of this class of compounds was significantly shorter than the isoxazole series.

**Figure 2:**
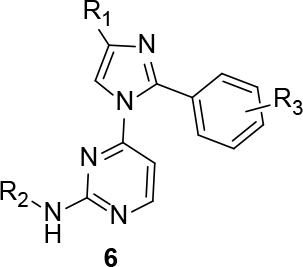
Imidazole PfPKG scaf-fold

Based on previous experience, we chose to focus first on exploration of the amino substituent on the pyrimidine. The synthesis of this set of derivatives is outlined in scheme 1. Commercially available 4-methyl-2-phenyl imidazole was deprotonated with sodium hydride then treated with 4-chloro-2-methythiopyrimidine in dry DMF at 60-70°C to arylate the imidazole nitrogen, followed by oxone-mediated conversion to the sulfone. Displacement of the sulfone with a variety of diamines, Boc-deprotection and acylation with preferred carboxylic acids as described previously^6^ afforded the target amides 10a-h.

**Scheme 1:**
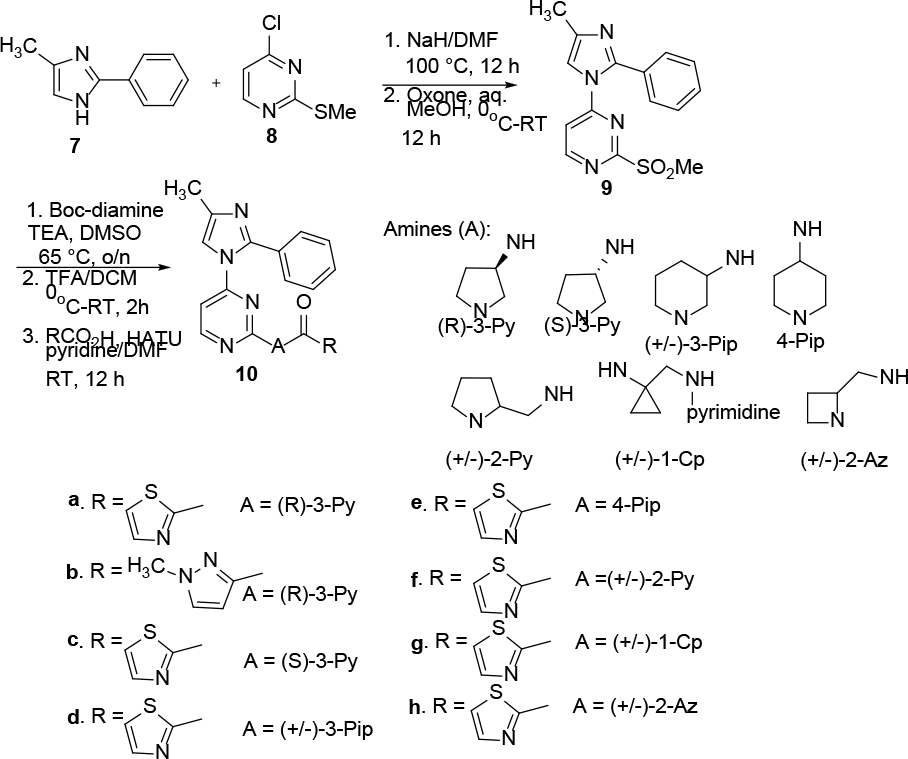
4-methyl imidazole PfPKG Inhibitors

It is evident from the data in table 3 that the (R)-3-aminopyrrolidine linker is strongly preferred compared to the other cyclic amine variations, a conformationally restricted alkyl diamine and the S-enantiomer of 3-aminopyrrolidine. Additionally, in this imidazole scaffold the potency difference between the 2-thiazolyl and 1-N-methyl-3-pyrazolyl amides is more pronounced, with a strong preference for the former. In this group, the most potent derivative (10a, PfPKG IC_50_ 320 nM) underwent further evaluation to provide baseline data for this chemotype on selectivity for human PKG and the T618Q mutant PfPKG as well as *in vitro* ADME characterization. Encouragingly, we observed excellent selectivity versus the related PKGs (3-10% inhibition @ 10 μM). *In vitro* ADME data revealed good water solubility, some inhibition of CYP3A4 and poor metabolic stability (table 2).

**Table 3:** *in vitro* PfPKG inhibition 4-methyl imidazole compounds **10a-10h**.

| Cpd | IC<sub>50</sub><br>(nM)/%<br>inh @ 1<br>μM | Cpd | IC<sub>50</sub><br>(nM)/%<br>inh @ 1<br>μM |
| --- | --- | --- | --- |
| **10a** | 320 | **10e** | 5% |
| **10b** | 1000 | **10f** | 3% |
| **10c** | 4% | **10g** | 6% |
| **10d** | 4% | **10h** | 2% |

We hypothesized that the 4-methyl group and unsubstituted aromatic ring were potential site(s) of metabolism. We chose to explore options at each site using low molecular weight changes. The 4-methyl group was replaced with a cyclopropyl ring and in view of existing isoxazole SAR on phenyl substituents, chose to target an unsubstituted and 3-chlorophenyl derivative. The synthesis, outlined in scheme 2, condensed appropriate chlorobenzamidines with cyclopropyl bromomethyl ketone to afford 4-cyclopropyl-2-phenyl imidazoles 13a and 13b in good yield. Following the steps outlined in scheme 1, the targets 14a and 14b were obtained in a straightforward manner. In parallel, using the 4-methylimidazole template, we explored a sampling of aryl substituents on the benzene ring to explore this feature of SAR. The synthesis of these analogs was accomplished as shown in scheme 3. Suzuki coupling between an appropriate boronic acid and commercially available 2-bromo-4-methyl imidazole afforded the corresponding 2-phenyl derivatives 17a-d that were processed as described in scheme 1 to afford the respective 2- and 4-chlorophenyl thiazolyl analogs 18b and 18c, respectively as well as the 3-methoxy and 3-trifluoromethoxy targets 18a and 18d. In both sets of analogs, we elected to use only the optimal 2-thiazolyl amide to provide the best comparison for activity versus 10a-h.

**Scheme 2:**
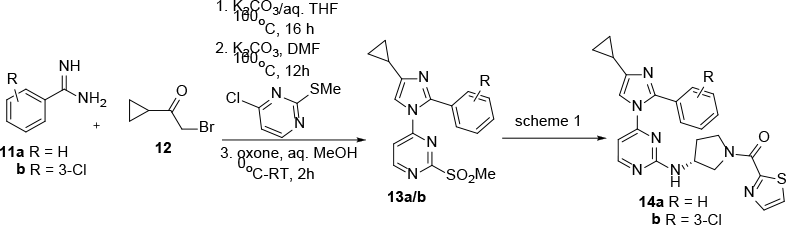
4-Cyclopropyl PfPKG Inhibitors

**Scheme 3:**
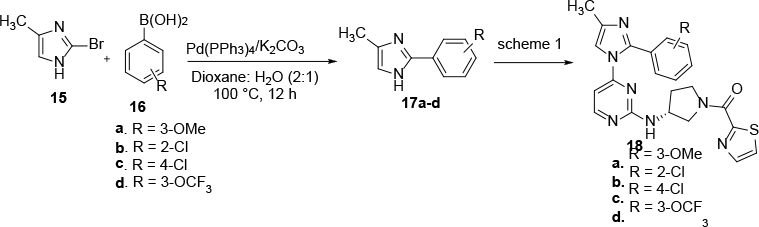
Phenyl-substituted PfPKG Inhibitors

Evaluation of these compounds as PfPKG inhibitors revealed that the cyclopropyl group in 14a provides a three-fold improvement in potency compared to 10a (Table 4) and that 3-chloro substitution provides a small additional benefit in 14b. Among the group of phenyl substituents evaluated, 3-chloro is preferred to its regioisomers 18b and 18c, similar to our previous observations. The other 3-substituted derivatives examined in this small set are comparable (18a, 18d) to 14a or less potent (18b, 18c). Collectively this structure-activity data suggests distinct SAR features compared to the isoxazole series.

**Table 4:** Cyclopropyl and aryl substituted PfPKG inhibitors

| CPD | IC<sub>50</sub><br>(nM)/%<br>inhib @<br>1 μM |
| --- | --- |
| 14a | 100 |
| 14b | 60 |
| 18a | 86 |
| 18b | 47% |
| 18c | 1160 |
| 18d | 150 |

The most potent example in this set, 14b, was examined for inhibition of human PKG and *P. falciparum* mutant T618Q, and as observed previously, demonstrated excellent selectivity against these two kinases with no inhibition at 10 μM.

These results led us to examine 14a and 14b in more detail with a focus on cellular activity and *in vitro* ADME. These imidazole-based PfPKG inhibitors have moderate to good water solubility, and show increased inhibition of CYP3A4 relative to the isoxazole series (table 2). Like the isoxazoles, there is poor metabolic stability and no measurable hERG inhibition at 10μM.

Using the HepG2 *P. berghei* sporozoite infectivity assay and 5 as a positive control, we chose to evaluate 10a and 14b to investigate a correlation between enzymatic and cellular activity. We were encouraged by the strong activity displayed by both compounds. The more efficacious imidazole, 14b, like 5, does not show a dose response (>90% at 2 and 10 μM), indicating that maximal inhibition was obtained at 2 μM. (Table 5). In this screen, 14b exhibited comparable activity to 5, representing a clear improvement compared to the isoxazoles 3 and 4. Although it is not definite from the limited concentrations employed in this assay, 14b appears to be more efficacious compared to the 10a, suggesting a potential correlation between *in vitro* enzymatic and cellular potency.

**Table 5:** Imidazole *P. berghei* infectivity assay

| Cpd | % decr<br>infn ±<br>SD @ 2<br>μM | % decr<br>infn ±<br>SD @ 10<br>μM | P value |
| --- | --- | --- | --- |
| 10a | 78±20 | 89±7 | < 0.001 |
| 14b | 94±3 | 93±3 | < 0.001 |
| 5 | 91±8 | 94±4 | < 0.001 |

This encouraging data led us to examine these two compounds against additional *Plasmodium* species in cell-based assays to more completely characterize this series and provide a baseline for future work. Imidazoles 10a and 14b were investigated in a dose-response assay that evaluated *P. cynomolgi* infectivity in prophylactic and radical cure modes. The prophylactic assay is a measure of the ability of *P. cynomolgi* to invade hepatocytes at the schizont stage. The radical cure mode evaluates activity against late stage or developing schizont stage parasites. The data in table 6 show that 14b is more active than 10a in the prophylactic mode with a sub-micromolar IC_50_, and interestingly has a modest effect in the radical cure mode. This data is consistent with a dose response effect for both compounds and indicates 14b is more efficacious than 10a in cell-based antiparasitic assays. Efficacy in the schizont model is important because there is a lower parasite burden at this stage compared to blood stage infection. Reduced sporozoite infectivity is known to reduce severity and infectivity.^9^ The data in table 5 also show these two compounds, unlike the controls, are comparatively non-toxic to host cells.

**Table 6:** *P. cynomolgi* infectivity cell-based assays

| Compound | Prophylactic Mode |  |  |
| --- | --- | --- | --- |
| | Schizont<br>IC<sub>50</sub><br>(μM) | Hypnozoite<br>IC<sub>50</sub><br>(μM) | Toxicity<br>(μM) |
| 14b | 0.93 | 9.88 | >20 |
| 10a | 7.91 | > 20 | >20 |
| Maduramicin | 0.01 | 0.02 | 5.78 |
| Tafenoquine | 0.19 | 0.14 | 13.82 |
| Compound | Radical Cure Mode |  |  |
| | Schizont<br>IC<sub>50</sub><br>(μM) | Hypnozoite<br>IC<sub>50</sub><br>(μM) | Toxicity<br>(μM) |
| 14b | 4.62 | > 20 | > 20 |
| 10a | 11.59 | > 20 | > 20 |
| Maduramicin | 0.03 | 0.03 | 11.91 |
| Tafenoquine | 3.61 | 3.50 | 18.61 |

When comparing these new PfPKG inhibitors to known inhibitors (figure 3), it is not surprising that enzymatic potency, while important, is not the sole determinant for cellular activity. It is encouraging to note the comparable enzymatic activity of 5 (IC_50_ 20 nM)^4^ and 14b (IC_50_ 60 nM) resulted in similar cellular activity and that a less potent imidazole analog 10a is measurably less efficacious. One observation based on data in this paper is that molecular weight contributes, based on a comparison of 3, 4, 14b and 5 in which the two highest molecular weight compounds show at best weak activity in cells. We have shown that like 5, the two isoxazoles are competitive PfPKG inhibitors that bind in the ATP pocket of the enzyme^9^. Studies are ongoing with imidazoles such as 14b to determine to mode of action for this class of compounds, along with investigations into obtaining what would be the initial report of an inhibitor bound to *P. falciparum* PKG.

**Figure 3:**
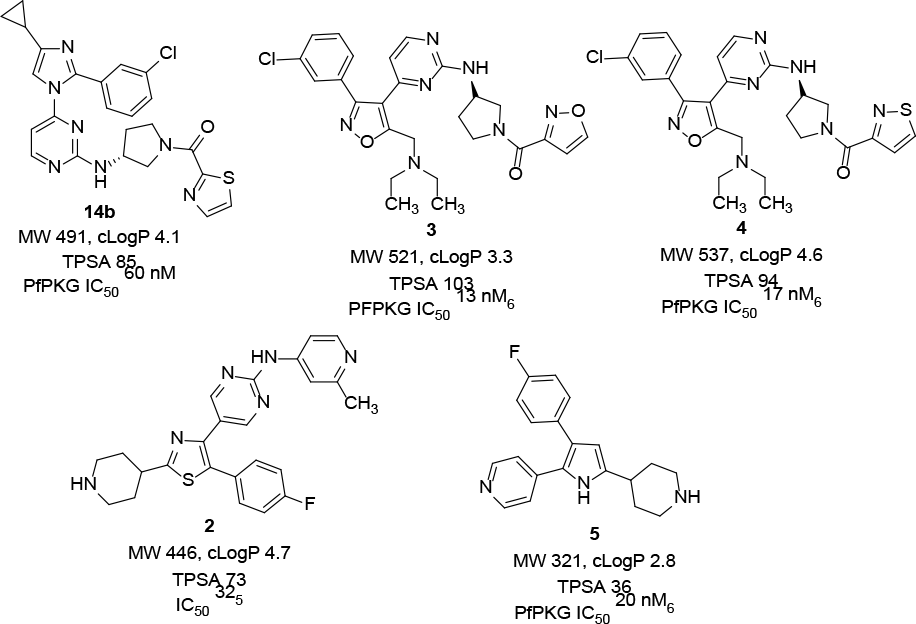
Chemical Property Comparison

It was gratifying to observe that like 3 and 4, neither of these imidazoles showed measurable hERG binding activity. This excitement was offset by the poor metabolic stability exhibited by both 14a and 14b, with half lives in human and mouse liver microsomes both less than 2 minutes (Table 2). In addition, an increase in CYP3A4 inhibition to less than 300 nM was recorded with both molecules and a decrease in water solubility, particularly in the case of 14b. Solutions to these weaknesses are under active investigation.

In conclusion, in response to the need to identify novel chemotypes as PfPKG inhibitors that lack the safety issues associated with many known scaffolds, we report the discovery of a new imidazole-based chemotype with enzymatic and cellular activity comparable to a literature standard, that lacks the hERG issues associated with other chemotypes and does not have any structural alerts associated with genotoxicity. Initial structure-activity relationships are distinct from known PfPKG inhibitors and while the ADME profile of lead 14b has weaknesses, the favorable profile in cellular infectivity assays and lack of general cytotoxic properties, provides the impetus to address the

ADME issues that currently exist. Those efforts are ongoing and will be reported in due course.

## Supporting information

biochemistry supplemental information

chemistry supplemental information

## ASSOCIATED CONTENT

### Supporting Information

Full experimental details on the synthesis and characterization of compounds, *in vitro* enzyme assay, cellular parasite infectivity and *in vitro* ADME assays are provided along with the manuscript in review cited as reference 7.

The Supporting Information is available free of charge on the ACS Publications website.

Chemistry-synthesis and characterization PDF

Biology: *in vitro* PfPKG assays, cellular parasite infectivity, *in vitro* ADME PDF

## AUTHOR INFORMATION

### Author Contributions

All authors have given approval to the final version of the manuscript.

## ACKNOWLEDGMENT

We acknowledge the Entomology Branch and Veterinary Medicine Branch AFRIMS, with special thanks to Ratawan Ubalee and team for the production of *P. cynomolgi*-infected mosquitoes.

## ABBREVIATIONS

PfPKG: *Plasmodium falciparum*
cGMP: dependent protein kinase
ATP: adenosine triphosphate
ADME: absorption, distribution, metabolism, elimination
CYP3A4: cytochrome P450 3A4
hERG: human ether-a-go-go related gene

---

Authors are required to submit a graphic entry for the Table of Contents (TOC) that, in conjunction with the manuscript title, should give the reader a representative idea of one of the following: A key structure, reaction, equation, concept, or theorem, etc., that is discussed in the manuscript. Consult the journal’s Instructions for Authors for TOC graphic specifications.

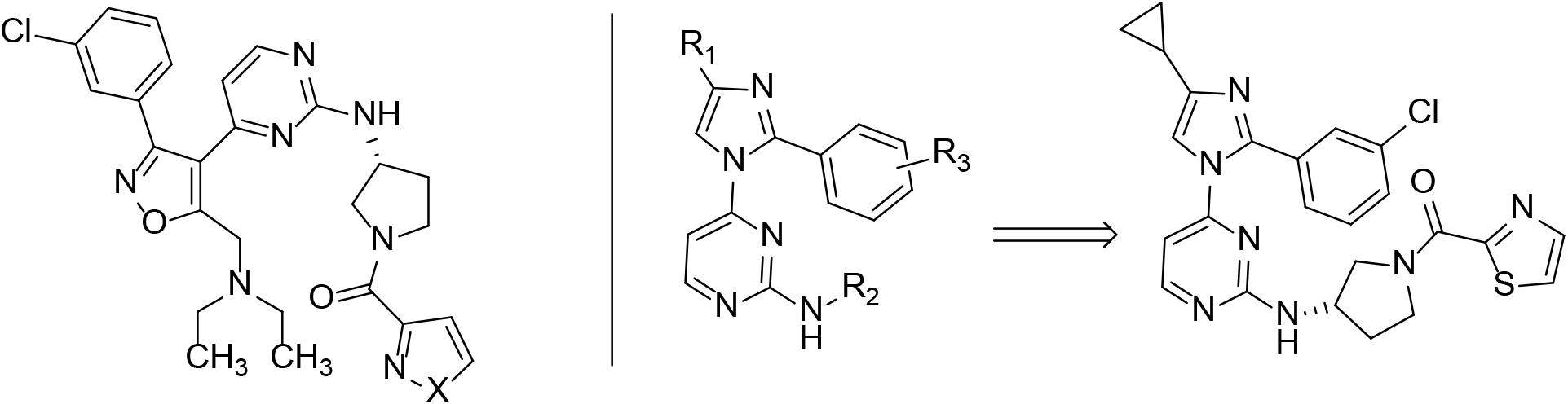

## REFERENCES

1. Arendse, L. B.; Wyllie, S.; Chibale, K.; Gilbert, I. H. Plasmodium kinases as potential drug targets for malaria: challenges and opportunities. ACS Infect. Dis. 2021, 7, 518–534.

2. Vanaerschot, M.; Murithi, J. M.; Pasaje, C.; Ghidelli-Disse, S.; Dwomoh, L.; Bird, M.; Spottiswoode, N.; Mittal, N.; Arendse, L. B.; Owen, E. S.; Wicht, K. J.; Siciliano, G.; Bösche, M.; Yeo, T.; Kumar, S. T. R.; Mok, S.; Carpenter, E.; Giddins, M. J.; Sanz, O.; Ottilie, S.; Alano, P.; Chibale, K.; Llinás, M.; Uhlemann, A. C.; Delves, M.; Tobin, A.; Doerig, C.; Winzeler, E.; Lee, M. C. S.; Niles, J.; Fidock, D. A. Inhibition of the resistance-refractory P. falciparum kinase PKG delivers prophylactic, blood stage and transmission-blocking antiplasmodial activity. Cell Chem. Biol. 2020, 27, 806–816.

3. Baker, D.A.; Stewart, L.B.; Large, J.M.; Boyer, P.W.; Ansell, K.H.; Jiménez, M.B.; El Bakkouri, M.; Birchall, K.; Dechering, K.J.; Bouloc, N.S.; Coombs, P.J.; Whalley, D.; Harding, D.J.; Smiljanic-Hurley, E.; Weldon, M.C.; Walker, E.M.; Dessens, J.T.; Lafuente, M.J.; Sanz, L.M; Gamo, F.-J.; Ferrer, S.B.; Hui, R.; Bousema, T.; Angulo-Barturén, I.; Merritt, A.T.; Croft, S.L.; Gutteridge, W.E.; Kettleborough, C.A.; Osborne, S.A.; A potent series targeting the malarial cGMP-dependent protein kinase clears infection and blocks transmission, Nat. Commun. 2017, 8, 430–439.

4. Penzo, M.; de las Heras-Dueña, L.; Mata-Cantero, L.; Diaz-Hernandez, B.; Vazquez-Muñiz, M.-J.; Ghidelli-Disse, S.; Drewes, G.; Fernández-Álvaro, E.; Baker, D.A., High-throughput screening of the Plasmodium falciparum cGMP-dependent protein kinase identified a thiazole scaffold which kills erythrocytic and sexual stage parasites, Sci. Reports, 2019, 9, 7005–7018.

5. Matralis, A.N.; Malik, A.; Penzo, M.; Moreo, I.; Almela, M.J.; Camino, I.; Crespo, B.; Saadeddin, A.; Ghidelli-Disse, S.; Rueda, L.; Calderon, F.; Osborne, S.A.; Drewes, G.; Böesche, M.; Fernández-Álvaro, E; Hernando, J.I.M.; Baker, D.A. Development of chemical entities endowed with potent, fast-killing properties against Plasmodium falciparum malaria parasites, J. Med. Chem. 2019, 62, 9217–9235.

6. Mahmood, S.U.; Cheng, H.; Tummalapalli, S.R.; Chakrasali, R.; Bheemanaboina, R.R.Y.; Kreiss, T.; Chojnowski, A.; Eck. T.; Siekierka, J.J.; Rotella, D.P. Discovery of isoxazolyl-based inhibitors of Plasmodium falciparum cGMP-dependent protein kinase, RSC Med. Chem. 2020, 11, 98–101.

7. Biftu, T.; Feng, D.; Ponpipom, M.; Girotra, N.; Liang, G.-B.; Qian, X.; Bugianesi, R.; Simeone, J.; Chang, L.; Gurnett, A.; Liberator, P.; Dulski, P.; Leavitt, P.S.; Crumley, T.; Misura, A.; Murphy, T.; Rattray, S.; Samaras, S.; Tamas, T.; Mathew, J.; Brown, C.; Thompson, D.; Schmatz, D.; Fisher, M.; Wyvratt, M.; Synthesis and SAR of 2,3-diarylpyrrole inhibitors of parasite cGMP-dependent protein kinase as novel anticoccidial agents, Bioorg. Med. Chem. Lett. 2005, 15, 3296–3301.

8. Govindasamy, K.; Jebiwott, S.; Jaiyan, D.K.; Davidow, A.; Ojo, K.K.; Van Voorhis, W.C.; Brochet, M.; Billker, O.; Bhanot, P., Invasion of hepatocytes by sporozoites requires cGMP-dependent protein kinase and calcium dependent protein kinase 4, Mol. Microbiol. 2016, 102, 349–363.

9. Alonso, P.L.; Sacarlal, J.; Aponte, J.J.; Leach, A.; Macete, E.; Aide, P.; Sigauque, B.; Milman, J.; Mandomando, I.; Bassat, Q.; Guinovart, C.; Espasa, M.; Corachan, S.; Lievens, M.; Navia, M.M.; Dubois, M.C.; Menendez, C.; Dubovsky, F.; Cohen J.; Thompson, R.; Ballou, W.R., Duration of protection with RTS,S/ASO2A malaria vaccine in prevention of Plasmodium falciparum disease in Mozambican children: single-blind extended follow up of a randomized controlled trial, Lancet, 2005, 366, 2012–2018.

10. Manuscript under review-see supplementary information.

