## Supplementary material for "Discovery of novel potent, cell-permeable inhibitors of *P. falciparum* cGMP-dependent protein kinase": biochemistry supplemental information

Biochemistry and Biology Supporting Information

+Sokol Institute for Pharmaceutical Life Sciences and Department of Chemistry and Biochemistry, Montclair State University, Montclair, New Jersey 07043, United States; %-Moulder Center for Drug Discovery Research, Temple University, Philadelphia PA, 19140; $-Department of Microbiology, Biochemistry and Molecular Genetics, Rutgers New Jersey Medical School, Newark NJ 07103; #-Department of Drug Discovery. Experimental Therapeutics Branch, Walter Reed Army Institute of Research, 503 Robert Grant Avenue, Silver Spring MD 20910.

***

**PfPKG Enzymatic Assays**

**IC­_50_ determination**

IC_50_ values were determined using the commercial immobilized metal ion affinity-based fluorescence polarization (IMAP) assay (Molecular Devices # R8127). In a 20 µL reaction, the following amounts of enzymes were used in each assay well: 17 ng of wild type PfPKG, 7 ng of T618Q PfPKG, and 10 ng of hPKG. Enzymes were preincubated at room temperature for 15 minutes with inhibitor concentrations ranging from 4 nM to 10 µM in assay buffer RB-T (10 mM Tris-HCl, pH 7.2, 10 mM MgCl_2_, 0.05% NaN_3_, 0.01% Tween®20). Next, 120 nM fluorescent peptide substrate (FAM-PKAtide or FAM-IP3R-derived peptide), 10 μM ATP, 1 μM cGMP, and 1.0 mM DTT were added to each well to begin the reaction. After a 60-minute incubation at room temperature, the reaction was halted and developed with 60 µL of the progressive binding reagent (PBR). For PfPKG and the mutant enzyme, the PBR was diluted 400-fold in 100% 1X IMAP Progressive Binding Buffer A and incubated for 30 minutes. For the human PKG, the PBR was diluted 400-fold in 75% 1X IMAP Progressive Binding Buffer A and 25% 1X IMAP Progressive Binding Buffer B, then incubated for 60 minutes. Fluorescent polarization was read in parallel and perpendicular to the excitation plane (ex. 485 nm/ em. 528 nm) using a Synergy 2 Microplate reader (BioTek, Winooski, VT). Two replicate wells were used for each sample and averaged before further analysis. The data were analyzed using a four-parameter logistic curve using Microsoft Excel Solver to make dose response curves in Microsoft Excel. To compare potency and assay quality, **5** was used as a positive control in each experiment.

**In vitro ADME**

Metabolic stability: Incubate samples at 37^o^C in the presence of human or mouse liver microsomes and NADPH according to standard methods (1). Aliquots are removed at 5 time points, quenched and analyzed (LCMS/MS and MS/MS as needed) for remaining test compound, along with a positive control. Microsomal protein content is adjusted to give consistent results. Data are reported as half-life and clearance. Assay acceptance criteria is 20% for all standards and 25% for the LLOQ.

Aqueous solubility: Kinetic aqueous solubility is measured by adding ~2 mg samples to pH 7.4 buffered aqueous solutions at 25^o^C. The mixture is agitated at 25^o^C for 1 hour, filtered and evaluated by UV and/or LCMSMS analysis. Data are reported as mg/mL and μM concentration.

CYP inhibition: Compounds are assessed for their ability to inhibit the three major human cytochrome P450 enzymes, 3A4, 2D6 and 2C9. Expressed enzymes are used to minimize non-specific binding and membrane partitioning issues (2). The 3A4 assay uses testosterone as a substrate and is analyzed by LC/MS/MS on a Waters TQ instrument using positive or negative electrospray ionization. The 2D6 and 2C9 assays use fluorescent substrates and are analyzed on an Envision Plate Reader.

hERG inhibition: HEK293 cells stably transfected with the hERG ion channel are grown to 80% confluency and then seeded into poly-lysine coated plates (25,000 cells/well). Cells are loaded with Thallos dye, treated with test compounds at 1 and 10 µM (final DMSO concentration = 0.1%) and then thalium flux is measured on a plate reader (excitation @ 480 nM; emission @ 530 nM) according to published procedures (3,4). Data are reported as percent inhibition at both concentrations.

**Cellular parasite infectivity assays:**

***In vitro P. cynomolgi* assay**

The *P. cynomolgi* liver stage experiment was performed as previously described^5^. Briefly, cryopreserved primary non-human primate hepatocytes (lot KXA) and hepatocyte culture medium (HCM) (InVitroGroTM CP Medium) were obtained from BioIVT, Inc., (Baltimore, MD, USA) and thawed following manufacturer recommendations. The hepatocytes were plated into pre-collagen coated 384-well plates (Cat No. 781956, Greiner, Monroe, NC, U.S.A.) and used for experiments within 2 – 4 days after plating. Infectious sporozoites were obtained from *An. dirus mosquitoes* infected with *P. cynomolgi* B strain and diluted accordingly (600 sporozoites/µL) in HCM. Compounds were dissolved in 100% DMSO and used at a final starting concentration of 20 µM in a 12-point, 3-fold serial dilution. The compounds were dosed in two treatment modes, prophylactic and radical cure. In prophylactic mode, drug was present for 4 days starting at point of sporozoite addition. Alternatively, in radical cure mode, drug was present for 4 days starting on day 4 post sporozoite inoculation.

Imaging and data analysis of the drug plates were completed using the Operetta CLS Imaging System and Harmony software 4.16 (Perkin Elmer, Waltham, MA, USA). Images were acquired using TRITC, DAPI, and bright field channels using a 10x objective. Following methodology previously described^6^, hepatocyte health (toxicity) and parasite populations (schizonts and hypnozoites) were identified and quantified by the specific object properties including area, mean intensity, maximum intensity, and cell roundness. Percent inhibition was calculated using the following equation, % inhibition = 100 x [(X – AVG N_ctrl_) / (AVG P_ctrl_ – AVG N_ctr_l)] where X is the replicate average at the specific compound concentration, AVG P_ctrl_ is the replicate average of the assay positive control (tafenoquine), and AVG Nctrl is the replicate average of the assay negative control (DMSO). The percent inhibition values were used to calculate the half maximal inhibitory concentration (IC_50_) and dose-response modeling was performed in GraphPad Prism version 8.1.0 (GraphPad, La Jolla, CA, USA) using the log(inhibitor) vs. response -- Variable slope (four parameters) model. The reported IC_50_ values are expressed as mean percent inhibition ± standard deviation (S.D.) from experimental replicates (n=2) with biological replicates (n=2) for both drug treatment modes.

***P. berghei* sporozoite infectivity assay:**

HepG2 cells were plated in 8-well LabTek chamber slides were infected with approximately 2 x 10^4^ luciferase-expressing *P. berghei* sporozoites in the presence of compounds or vehicle. Compound-containing media was replaced with compound-free medium at 16 post-infection. At 48 hours post-infection, medium was replaced with D-luciferin diluted in PBS. Luciferase activity was measured using an IVIS® Spectrum in vivo imaging system.

**References:**

1.	Lounsbury N, Mateo G, Jones B, Papaiahgari S, Thimmulappa RK, Teijaro C, Gordon J, Korzekwa K, Ye M, Allaway G, Abou-Gharbia M, Biswal S, Childers W, Jr. Heterocyclic chalcone activators of nuclear factor (erythroid-derived 2)-like 2 (Nrf2) with improved in vivo efficacy. Bioorg Med Chem. 2015;23(17):5352-9.

2.	McMasters DR, Torres RA, Crathern SJ, Dooney DL, Nachbar RB, Sheridan RP, Korzekwa KR. Inhibition of recombinant cytochrome P450 isoforms 2D6 and 2C9 by diverse drug-like molecules. Journal of medicinal chemistry. 2007;50(14):3205-13.

3.	Huang XP, Mangano T, Hufeisen S, Setola V, Roth BL. Identification of human Ether-a-go-go related gene modulators by three screening platforms in an academic drug-discovery setting. Assay Drug Dev Technol. 2010;8(6):727-42.

4.	Titus SA, Beacham D, Shahane SA, Southall N, Xia M, Huang R, Hooten E, Zhao Y, Shou L, Austin CP, Zheng W. A new homogeneous high-throughput screening assay for profiling compound activity on the human ether-a-go-go-related gene channel. Anal Biochem. 2009;394(1):30-8.

5.	Vanachayangkul, P.; Im-Erbsin, R.; Tungtaeng, A.; Kodchakorn, C.; Roth, A.; Adams, J.; Chaisatit, C.; Saingam, P.; Sciotti, R. J.; Reichard, G. A.; Nolan, C. K.; Pybus, B. S.; Black, C. C.; Lugo-Roman, L. A.; Wegner, M. D.; Smith, P. L.; Wojnarski, M.; Vesely, B. A.; Kobylinski, K. C., Safety, Pharmacokinetics, and Activity of High-Dose Ivermectin and Chloroquine against the Liver Stage of Plasmodium cynomolgi Infection in Rhesus Macaques. *Antimicrob Agents Chemother* **2020,** *64* (9).

6.	Roth, A.; Maher, S. P.; Conway, A. J.; Ubalee, R.; Chaumeau, V.; Andolina, C.; Kaba, S. A.; Vantaux, A.; Bakowski, M. A.; Thomson-Luque, R.; Adapa, S. R.; Singh, N.; Barnes, S. J.; Cooper, C. A.; Rouillier, M.; McNamara, C. W.; Mikolajczak, S. A.; Sather, N.; Witkowski, B.; Campo, B.; Kappe, S. H. I.; Lanar, D. E.; Nosten, F.; Davidson, S.; Jiang, R. H. Y.; Kyle, D. E.; Adams, J. H., A comprehensive model for assessment of liver stage therapies targeting Plasmodium vivax and Plasmodium falciparum. *Nat Commun* **2018,** *9* (1), 1837.
