## Supplementary material for "Discovery of novel potent, cell-permeable inhibitors of *P. falciparum* cGMP-dependent protein kinase": chemistry supplemental information

Chemistry Supporting Information

+Sokol Institute for Pharmaceutical Life Sciences and Department of Chemistry and Biochemistry, Montclair State University, Montclair, New Jersey 07043, United States; %-Moulder Center for Drug Discovery Research, Temple University, Philadelphia PA, 19140; $-Department of Microbiology, Biochemistry and Molecular Genetics, Rutgers New Jersey Medical School, Newark NJ 07103; #-Department of Drug Discovery. Experimental Therapeutics Branch, Walter Reed Army Institute of Research, 503 Robert Grant Avenue, Silver Spring MD 20910.

***

**Data for the following chemical structures**

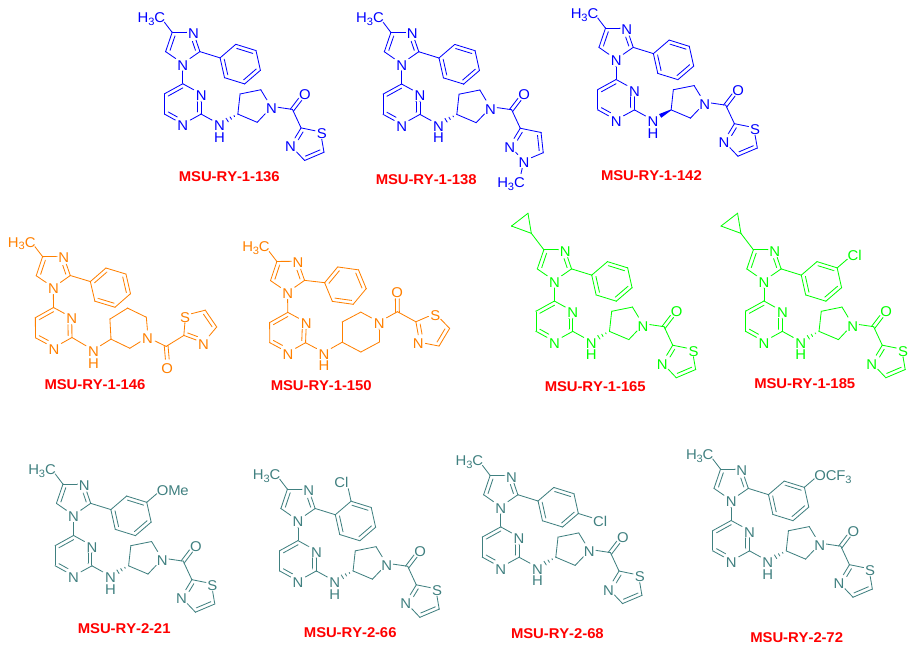

General chemical procedures:

Compound characterization: All reagents and solvents were used as received from
commercial suppliers. Compounds were analyzed using a UPLC system with a BEH-C18column (2.1 cm x 50 mm, 0-100% acetonitrile/water gradient over 6 minutes with UV 254 nmdetection) and an LCMS 2020 system (LC18 25 cm x 4.6 mm 5 μM column, 0-95%acetonitrile/water gradient over 10 minutes; UV monitor at 220 and 254 nM). Thin layerchromatography was done on silica gel G plates with UV or phosphomolybdic acid detection.Mass spectra were recorded on a CMS system with ESI probe. All of the reported yields are for isolated products and compounds were purified by automated flash chromatography. Proton NMR and ^13^C spectra were obtained at 400 and 101 MHz, respectively, in CDCl_3_ unless otherwise stated. All final compounds had purities of at least 95% based on 1H NMR and UPLC analyses.

**General route for synthesis of 10a-h:**

**Arylation:** To the solution of **7** (1 eq) in anhydrous DMF (10 ml) at ambient temperature was added NaH (1.2 eq, 60% in mineral oil) and the resulting mixture was stirred for 30 minutes. at 0°C followed by addition of 4-chloro-2-(methylthio)pyrimidine (1.2 eq). The resulting reaction mixture then was stirred for 12 h with temperature gradually rising from 0 °C to ambient temperature, then quenched by addition of water (10 ml). The quenched mixture was partitioned between ethyl acetate/H_2_O (3 times). Organic layer was collected, washed with brine, dried, concentrated and purified by CombiFlash silica gel chromatography eluting with a mixture of ethyl acetate/hexane to provide the desired product as an off-white solid.

**9:** Pyrimdinyl methyl sulfide was dissolved at room temperature in a mixture of methanol/distilled water (2:1 ratio, final concentration ~0.2M). Oxone (potassium peroxymonosulfate, 2.1 eq) was added at room temperature and the reaction was stirred overnight. The reaction mixture was poured into water and extracted with 4 portions of ethyl acetate. The collected organic extracts were washed with brine, dried, concentrated and purified by CombiFlash silica gel chromatography eluting with a mixture of ethyl acetate/hexane to provide the desired sulfone as a white solid.

**Amine displacement of sulfone**: **9** was dissolved in reagent grade DMSO (~0.3M) at room temperature. Triethylamine (3 equivalents) were added, followed by 2 equivalents of the desired Boc-protected diamine. The reaction was heated under nitrogen to 65 °C overnight. After cooling to room temperature the reaction mixture was poured into brine solution and extracted 4 times with ethyl acetate. The collected organic extracts were washed with brine, dried and concentrated. The desired product was isolated by CombiFlash silica gel chromatography eluting with 5-10% methanol in dichloromethane to furnish pure product.

**Boc Deprotection**: The respective Boc-amine (1 eq) was dissolved in 10 mL methylene chloride was added excess trifluoro acetic acid at 0 °C. The reaction mixture was stirred for 2 hours at room temperature. Solvent was removed in vacuo, the residue was dissolved in brine solution and quenched it with solid NaHCO3 till the pH is neutral and extracted 4 times with ethyl acetate. The collected organic extracts were washed with brine, dried and concentrated. The desired product was isolated by CombiFlash silica gel chromatography eluting with 10% methanol in dichloromethane to furnish pure product.

**Amide coupling**: To the solution of the appropriate amine in reagent grade DMF at room temperature was added 3 equivalents of pyridine, followed by 1.3 equivalents of HATU and 1.2 equivalents of the desired carboxylic acid. The reaction was stirred at room temperature overnight. In the morning, the reaction mixture was poured into 5 volumes of water and extracted with 4 portions of ethyl acetate. The collected organic extract was washed with two portions of brine, dried and concentrated. The product was purified by CombiFlash silica gel chromatography eluting with 5-10% methanol in dichloromethane to furnish the desired amine product.

Yields stated with each compound reflect this last reaction.

**10a** Yield: 62%; δ 8.13 (dd, *J* = 5.4, 1.6 Hz, 1H), 7.90 (dd, *J* = 7.4, 3.2 Hz, 1H), 7.54 (dd, *J* = 3.1, 1.4 Hz, 1H), 7.49 – 7.40 (m, 2H), 7.40 – 7.31 (m, 3H), 7.29 (d, *J* = 0.9 Hz, 1H), 7.10 – 6.72 (m, 5H), 6.17 (s, 1H), 5.50 (s, 1H, NH), 4.42 – 4.26 (m, 2H), 4.02 – 3.68 (m, 3H), 2.31 (s, 3H), 2.01 (s, 1H), 1.86 (dd, *J* = 14.5, 7.5 Hz, 1H); ^13^C δ 165.3, 165.2, 161.8, 159.7, 159.4, 159.1, 157.3, 146.5, 143.9, 143.8, 143.6, 143.6, 138.5, 129.0, 128.4, 124.2, 116.4, 116.3, 47.1, 45.7, 32.3, 29.5, 13.6. ESI-MS: m/z 432.0 [M + H]^+^.

**10b** Yield: 58%; white solid; δ- 8.39 (dd, *J* = 9.8, 6.4, 1H), 8.12 (dd, *J* = 5.4, 0.9 Hz, 1H), 7.37 – 7.26 (m, 6H), 6.77 (d, *J* = 2.3 Hz, 1H), 6.12 (s, 1H), 4.17 (s, 1H), 4.06 (dd, *J* = 14.1, 7.2 Hz, 1H), 3.91 – 3.84 (m, 4H), 3.77 (dd, *J* = 19.6, 13.7 Hz, 2H), 2.28 (s, 3H), 2.12 (s, 1H), 1.94 (s, 1H). ^13^C: δ 162.2, 161.8, 157.0, 147.5, 147.4, 140.3, 137.6, 137.5, 135.2, 129.0, 128.6, 128.4, 128.3, 109.1, 108.7, 46.9, 44.8, 22.7, 13.1. ESI-MS: m/z 429.0 [M + H]^+^.

**10c** Yield: 60%; white solid; δ 8.36 (dd, *J* = 10.4, 5.0 Hz, 1H), 8.15 – 7.78 (m, 1H), 7.91 – 7.87 (m, 1H), 7.54 – 7.26 (m, 6H), 6.49 (dd, *J* = 18.5, 4.5 Hz, 1H), 5.49 (s, 1H), 4.55 – 4.18 (m, 2H), 4.08 (dd, *J* = 12.5, 4.9 Hz, 1H), 3.67 (dd, *J* = 85.2, 36.0 Hz, 2H), 2.31 (s, 3H), 2.18 – 1.94 (m, 1H), 1.94 – 1.65 (m, 1H). ^13^C: δ 165.2, 162.1, 160.6, 160.3, 159.1, 134.2, 128.5, 128.3, 124.2, 124.1, 114.9, 114.8, 47.1, 45.7, 38.6, 12.0. ESI-MS: m/z 432.0 [M + H]^+^.

**10d** Yield: 53%; white solid; δ 8.35 (s, 1H), 7.89 (s, 1H), 7.68 (s, 1H), 7.54 (s, 1H), 7.47 – 7.31 (m, 5H), 6.47 (s, 1H), 6.22 (s, 1H), 4.19 (s, 2H), 4.06 – 3.73 (m, 1H), 3.48 – 3.36 (m, 2H), 2.33 (s, 3H), 1.74 (s, 2H), 1.59 (s, 2H); ^13^C: δ 162.2, 160.3, 143.1, 142.7, 132.0, 129.1, 128.6, 128.4, 128.3, 127.6, 127.4, 124.1, 123.7, 50.5, 47.7, 44.2, 34.7, 31.6, 25.3, 22.6, 11.9; ESI-MS: m/z 446.0 [M + H]^+^.

**10e** Yield: 57%; off-white solid; δ-8.37 (dd, *J* = 4.6, 2.6 Hz, 1H), 7.88 – 7.85 (m, 1H), 7.52 – 7.51 (m, 1H), 7.36 – 7.26 (m, 6H), 6.55 (s, 1H), 5.43 – 5.26 (m, 2H), 4.49 (s, 1H), 3.65 (s, 1H), 3.18 (s, 1H), 2.81 – 2.79 (m, 1H), 2.30 (s, 3H), 1.73 – 1.68 (m, 2H), 1.25 – 1.24 (m, 2H); ^13^C NMR: δ 165.1, 161.9, 161.8, 160.5, 160.3, 159.2, 143.2, 142.9, 134.3, 128.2, 124.1, 114.6, 48.3, 45.1, 42.6, 12.1; ESI-MS: m/z 446.0 [M + H]^+^.

**10f:** Yield 68%, white solid; *δ*= 8.3(d, 1H, H11, J=4.9Hz), 7.9(s, 1H, H21), 7.5(s, 1H, H20), 7.4-7.1(m, 7H, H2, H5, H6, H7, H5’, H6’, H10), 6.4(s, 1H, NH), 4.5-3.4(m, 5H, H13,H14, H15), 2.3 (s, 3H, H8), 2.3-1.6 (m, 4H, H15, H16). ESI (M+ H)^+^ +(MeOH + H)^+^ 480 m/z

**10g**: Yield 61%; white solid; *δ*-8.3(d, 1H, H11, *J*=4.8Hz), 7.8(s, 1H, H20), 7.7 (s, 1H, NH), 7.6(s, 1H, H19), 7.4-7.2(m,6H, H2, H5, H6, H7, H5’, H6’), 6.6 (s, 1H, NH), 6.4 (s,1H, H10), 3.4(s,2H, H13), 2.3 (s, 3H, H8), 1.0-0.8(m, 4H, H15,H16). ESI (M+ H)^+^ +(MeOH + H)^+^ 466 m/z

**10h**: Yield 71%; white solid;*δ*-8.3(d, 1H, H11, *J*=4.8Hz), 7.8(s, 1H, H20), 7.7 (s, 1H, NH), 7.6(s, 1H, H19), 7.4-7.2(m,6H, H2, H5, H6, H7, H5’, H6’), 6.6 (s, 1H, NH), 6.4 (s,1H, H10), 3.4(s,2H, H13), 2.3 (s, 3H, H8), 1.0-0.8(m, 4H, H15,H16). ESI (M+ H)^+^ +(MeOH + H)^+^ 466 m/z

**Synthesis of 14a/b**

**Preparation of cyclopropyl imidazole derivatives:** A mixture of benzamidine (1 eq), 2-bromo-1-cyclopropylethanone (1.2 eq) and potassium carbonate (2 eq) in 2:1 ratio of THF and water (50 ml) was stirred at reflux condition for 18 hours. After cooling to room temperature, the reaction mixture was filtered off and evaporated in vacuo. Then the residue was poured into water, extracted with ethyl acetate, dried over sodium sulfate, and evaporated in vacuo. The crude product was purified by CombiFlash silica gel chromatography eluting with 30-50% ethyl acetate in hexane to furnish the desired products.

The synthesis of **13a/b**, amine displacement, deprotection and acylation was carried out as described above. The yield stated below is for the acylation reaction to afford the target compound.

**14a.** Yield: 73%; off-white solid; δ 8.19 (d, *J* = 4.6 Hz, 1H), 7.95 (dd, *J* = 11.4, 3.2 Hz, 1H), 7.80 (d, *J* = 2.7 Hz, 1H), 7.41 – 7.34 (m, 6H), 6.44 (s, 1H), 4.16 (d, *J* = 5.6 Hz, 2H), 3.74 – 3.39 (m, 3H), 1.98 (s, 1H), 1.88 (dd, *J* = 9.6, 5.1 Hz, 2H), 0.89 – 0.75 (m, 4H); ^13^C: δ 165.4, 165.3, 161.7, 159.2, 159.1, 157.15, 144.5, 144.4, 129.2, 128.5, 124.2, 114.8, 114.6, 47.1, 45.6, 32.4, 29.6, 8.7, 7.2; ESI-MS: m/z 458.0 [M + H]^+^.

**14b.** Yield: 69%; off-white solid; δ 8.16 (d, *J* = 5.4, 2.4 Hz, 1H), 7.87 (dd, *J* = 4.9, 3.2 Hz, 1H), 7.52 – 7.48 (m, 2H), 7.41 – 7.09 (m, 4H), 6.18 (s, 1H), 4.29 (d, *J* = 5.3 Hz, 2H), 4.0 - 3.63 (m, 3H), 2.16 (s, 1H), 1.91 – 1.85 (m, 2H), 0.89 – 0.83 (m, 4H); ^13^C NMR (CDCl_3,_ 101 MHz): δ 165.3, 161.8, 159.1, 157.0, 145.0, 143.6, 134.3, 129.1, 124.2, 115.3, 115.1, 47.1, 45.6, 32.4, 29.6, 8.8, 7.1; ESI-MS: m/z 492.0 [M + H]^+^.

**Synthesis of 18a-d:**

**17a-d:** A solution of 2 M K_2_CO_3_ solution (3 ml) in dioxane (3 ml) was taken in round bottom flask and was purged with nitrogen balloon for 5 minutes at room temperature. A mixture of aryl boronic acid (**16a-d**, 1.2 eq) and 2-bromo 4-methyl imidazole (**15**, 1 eq) was added to this reaction mixture, and it was again purged with nitrogen for 5 min. Pd(PPh_3_)_4_ (0.05 eq) was then added, followed by purging and allowed the reaction mixture to stir at 90 °C for overnight. After completion of the reaction, product was extracted with ethyl acetate (2 × 50 ml). The combined organic layer was concentrated in vacuo and crude reaction mixture was purified by CombiFlash silica gel chromatography eluting with 30-50% ethyl acetate in hexane to furnish the desired products.

**17a-d** were processed as described above to obtain **18a-d**. The yield for each compound is for the acylation reaction to obtain the target compound.

**18a.** Yield: 71%; off-white solid; δ 8.10 (dd, *J* = 5.4, 2.7 Hz, 1H), 7.87 (dd, *J* = 7.4, 3.2 Hz, 1H), 7.51 (dd, *J* = 3.1, 2.6 Hz, 1H), 7.28 – 7.20 (m, 2H), 7.04 – 7.01 (m, 1H), 6.94 – 6.88 (m, 2H), 6.13 (s, 1H), 4.41 (s, 1H), 4.23 (s, 1H), 3.93 – 3.83 (m, 1H), 3.76 – 3.74 (m, 4H), 3.54 (dd, *J* = 4.8, 4.0 Hz, 1H), 2.28 (s, 3H), 2.04 (dd, *J* = 6.2, 5.6 Hz, 1H), 2.10 – 1.78 (m, 1H); ^13^C δ 165.3, 165.2, 161.8, 159.5, 157.0, 146.1, 143.6, 137.9, 129.5, 124.2, 124.1, 121.7, 121.4, 116.5, 116.4, 47.2, 45.7, 32.3, 29.5, 13.3; ESI-MS: m/z 462.0 [M + H]^+^.

**18b.** Yield: 71%; off-white solid; δ 8.17 (d, *J* = 5.5 Hz, 1H), 7.89 (dd, *J* = 10.4, 3.2 Hz, 1H), 7.54 – 7.53 (m, 2H), 7.37 (dd, *J* = 12.8, 6.0 Hz, 4H), 6.28 (s, 1H), 5.85 (s, 1H), 5.29 (s, 1H), 4.21 (s, 2H), 3.80 – 3.51 (m, 3H), 2.37 (s, 3H), 2.06 – 2.03 (m, 1H), 1.80 (s, 1H); ^13^C δ 165.2, 161.2, 159.1, 156.5, 143.9, 143.6, 133.9, 129.6, 127.1, 124.2, 115.1, 47.1, 45.6, 32.3, 29.5, 13.1; ESI-MS: m/z 466.0 [M + H]^+^.

**18c.** Yield: 71%; off-white solid; δ 8.25 (s, 1H), 7.92 – 7.89 (m, 1H), 7.55 – 7.46 (m, 3H), 7.40 – 7.29 (m, 3H), 6.26 (s, 1H), 4.29 (s, 3H), 3.86 – 3.80 (m, 3H), 2.39 (s, 3H), 2.07 (s, 1H), 1.85 (s, 1H); ^13^C δ 165.3, 161.8, 160.0, 159.2, 156.8, 143.7, 131.3, 131.2, 124.2, 116.8, 116.0, 47.1, 45.6, 32.3, 29.5, 12.8; ESI-MS: m/z 466.0 [M + H]^+^.

**18d.** Yield: 71%; off-white solid; δ 8.27 (d, *J* = 5.3, 3.1 Hz, 1H), 7.92 – 7.88 (m, 2H), 7.56 – 7.53 (m, 2H), 7.48– 7.46 (m, 1H), 7.36 – 7.31 (m, 2H), 6.29 (s, 1H), 4.26 (s, 2H), 4.08 (s, 1H), 3.93 – 3.59 (m, 2H), 2.38 (s, 3H), 2.08 (s, 1H), 1.95 (s, 1H); ^13^C δ 161.7, 159.7, 159.2, 157.4, 153.2, 149.9, 138.1, 129.0, 128.4, 119.7, 116.4, 46.8, 45.2, 32.3, 29.7, 13.3; ESI-MS: m/z 516.0 [M + H]^+^.

**^1^H, ^13^C and mass spectra of final compounds**

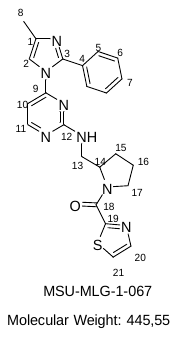

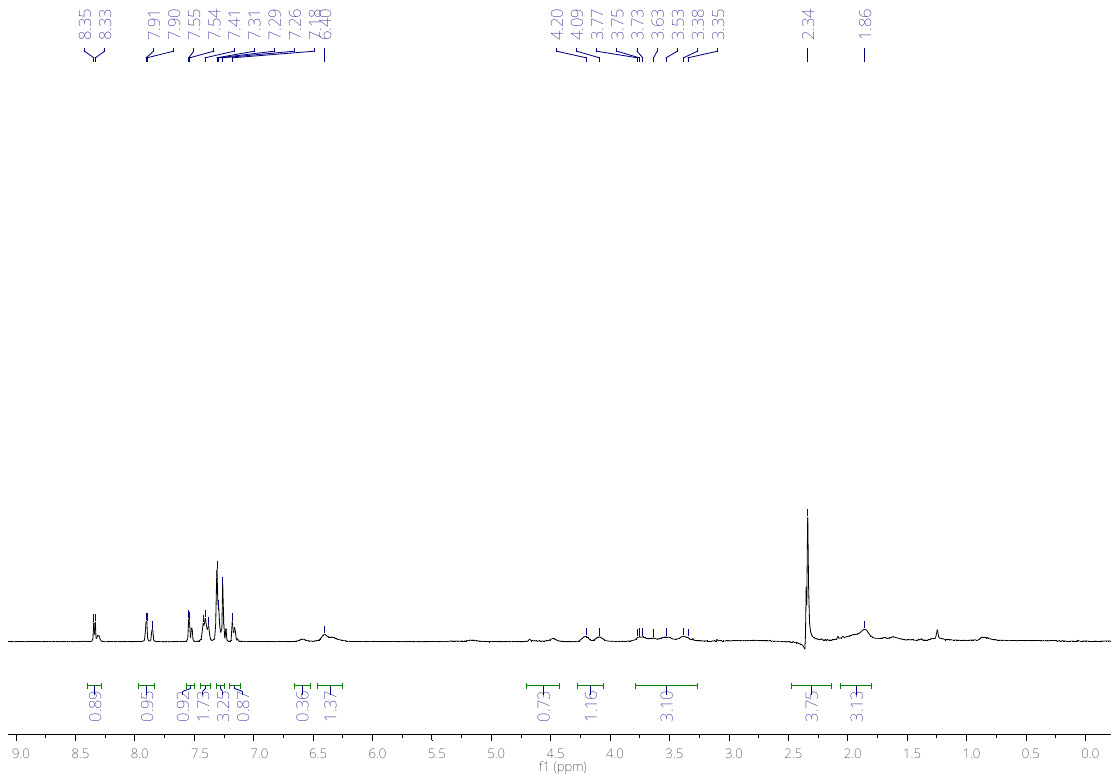

^1^H NMR (CDCl_3_, 400Hz) 10f

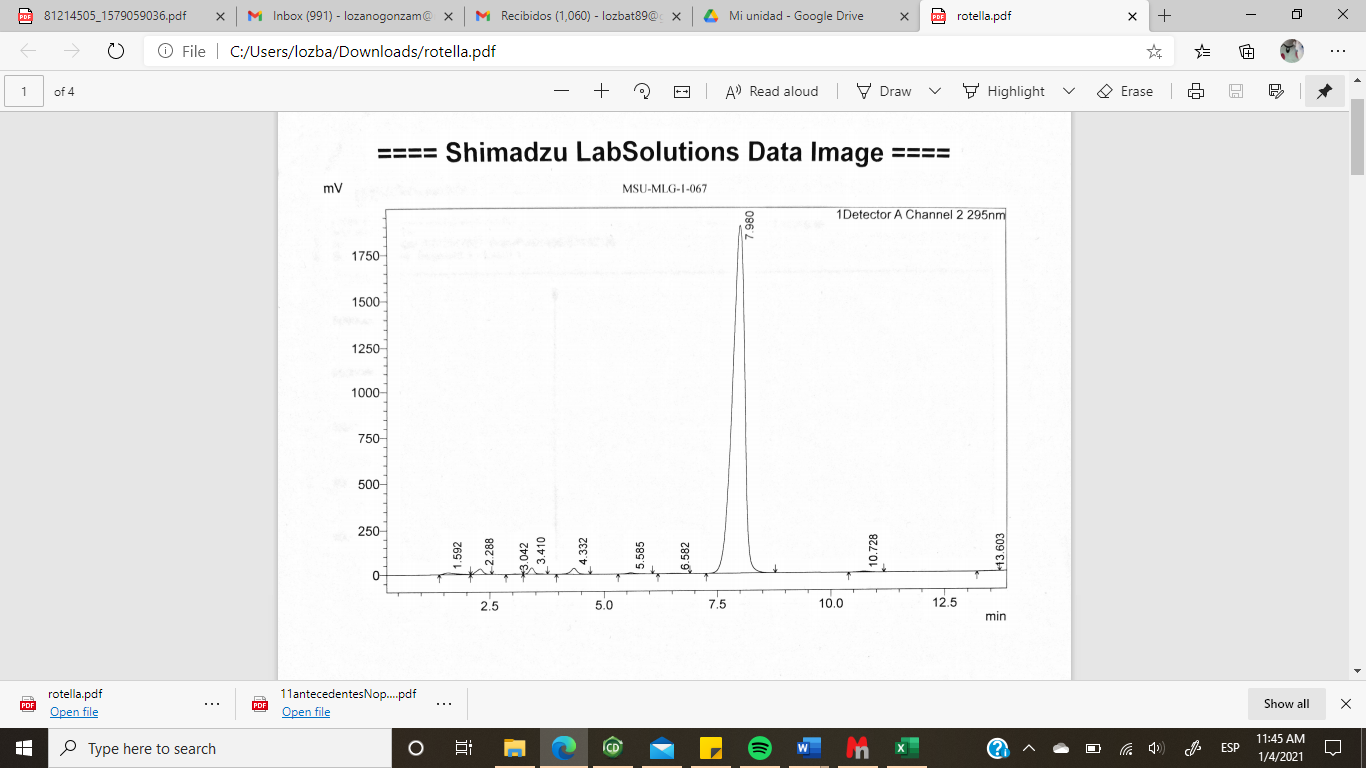

LC Spectrum 10f

**MS Spectrum** 10f (M+ H)^+^ +(MeOH + H)^+^ 480 m/z

**10g**

**^1^H NMR (400MHz, CDCl_3_)**: *δ*= 8.3(d, 1H, H11, *J*=4.8Hz), 7.8(s, 1H, H20), 7.7 (s, 1H, NH), 7.6(s, 1H, H19), 7.4-7.2(m,6H, H2, H5, H6, H7, H5’, H6’), 6.6 (s, 1H, NH), 6.4 (s,1H, H10), 3.4(s,2H, H13), 2.3 (s, 3H, H8), 1.0-0.8(m, 4H, H15,H16) ppm. **ME** (M+ H)^+^ +(MeOH + H)^+^ 466 m/z

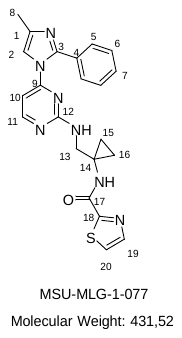

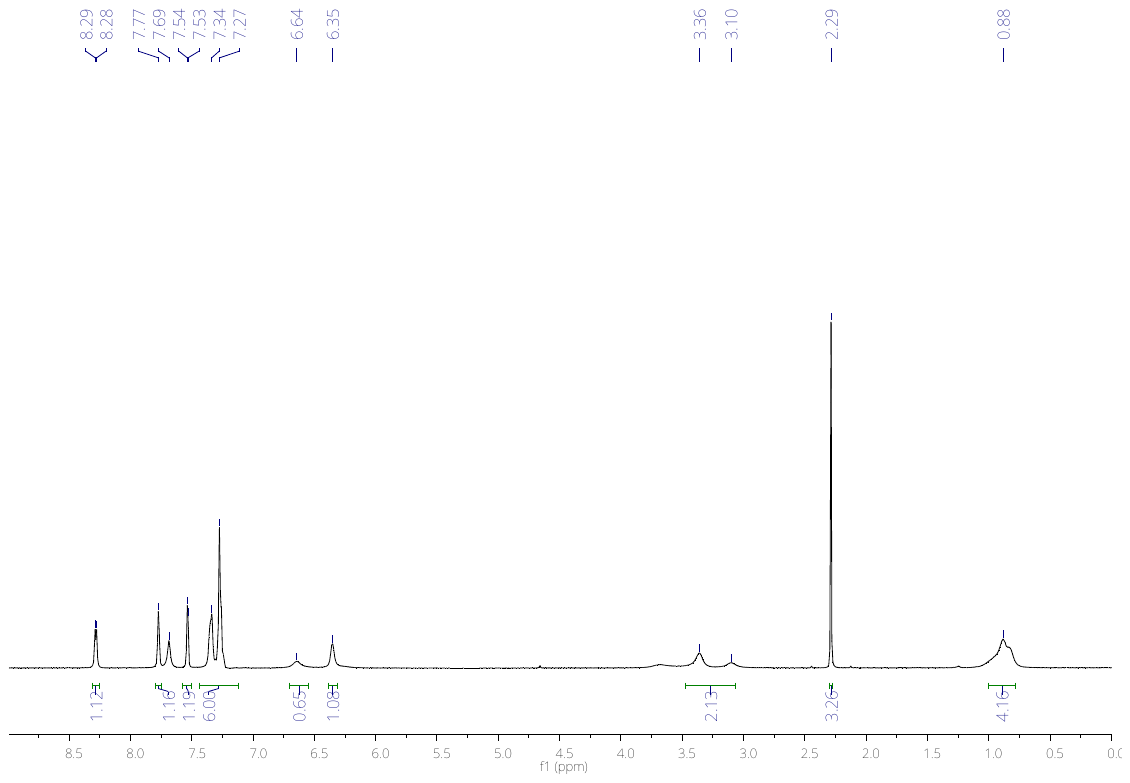

^1^H NMR (CDCl_3_, 400Hz) **10g**

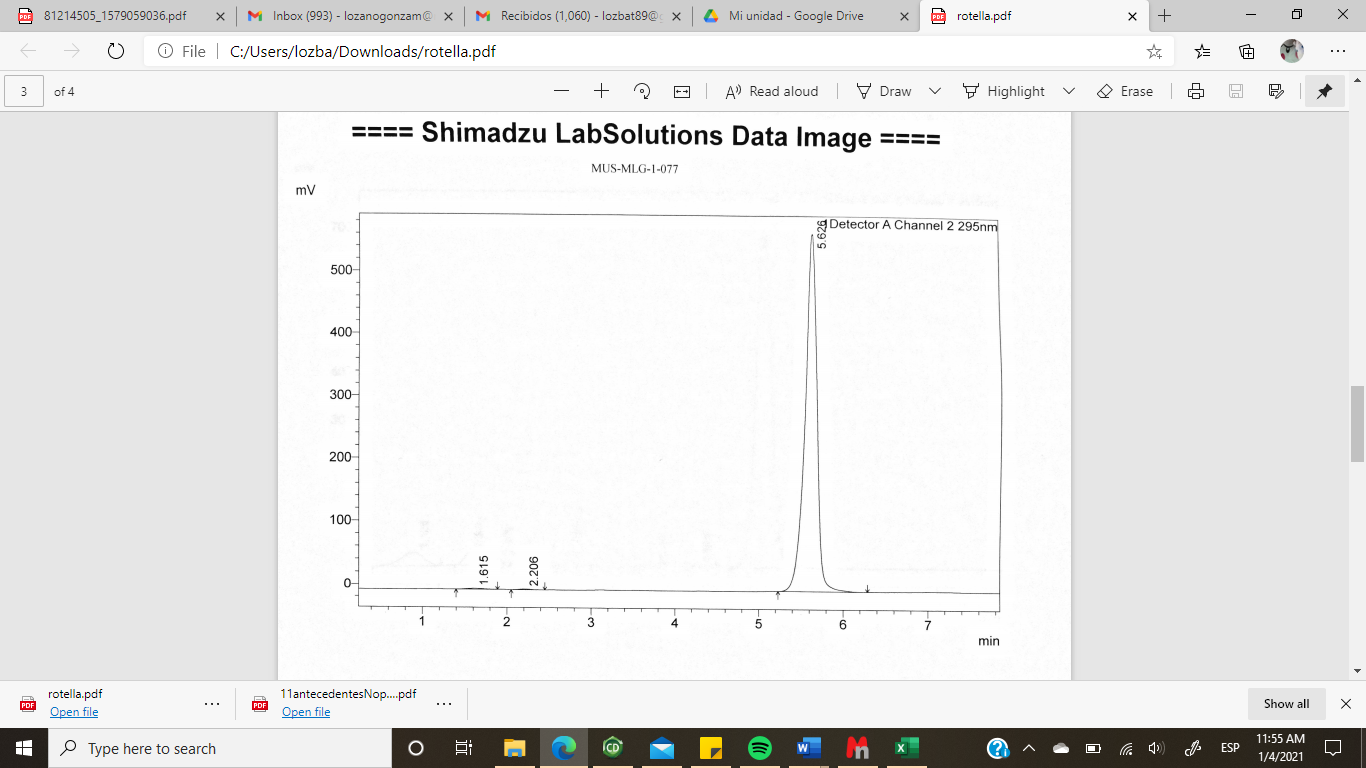

LC Spectrum **10g**

**MS Spectrum** **10g** (M+ H)^+^ +(MeOH + H)^+^ 466 m/z

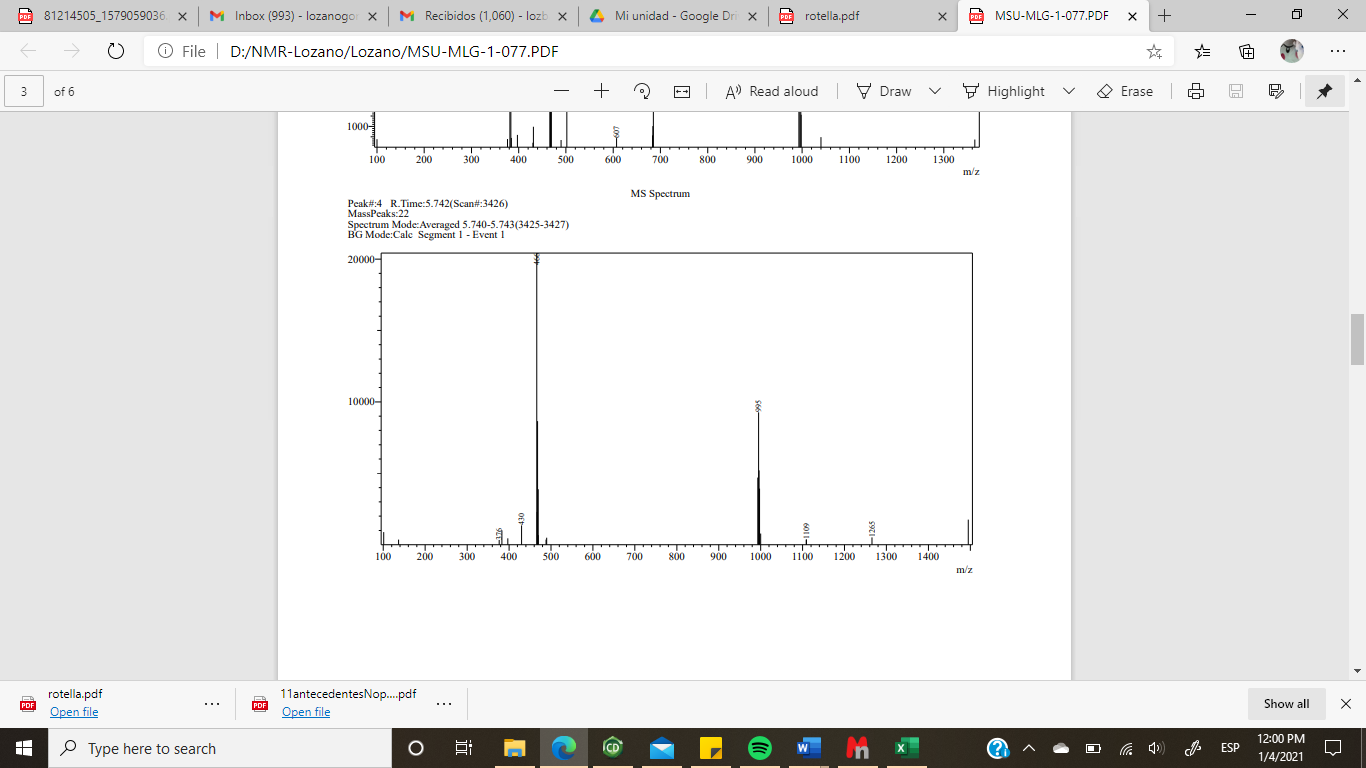

**10h**

^1^H NMR (400MHz, CDCl_3_): **= 8.3(d, 1H, H11, *J*=4.8Hz), 7.8(s, 1H, H20), 7.7 (s, 1H, NH), 7.6(s, 1H, H19), 7.4-7.2(m,6H, H2, H5, H6, H7, H5’, H6’), 6.6 (s, 1H, NH), 6.4 (s,1H, H10), 3.4(s,2H, H13), 2.3 (s, 3H, H8), 1.0-0.8(m, 4H, H15,H16) ppm. **ME** (M+ H)^+^ +(MeOH + H)^+^ 466 m/z

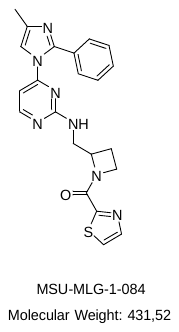

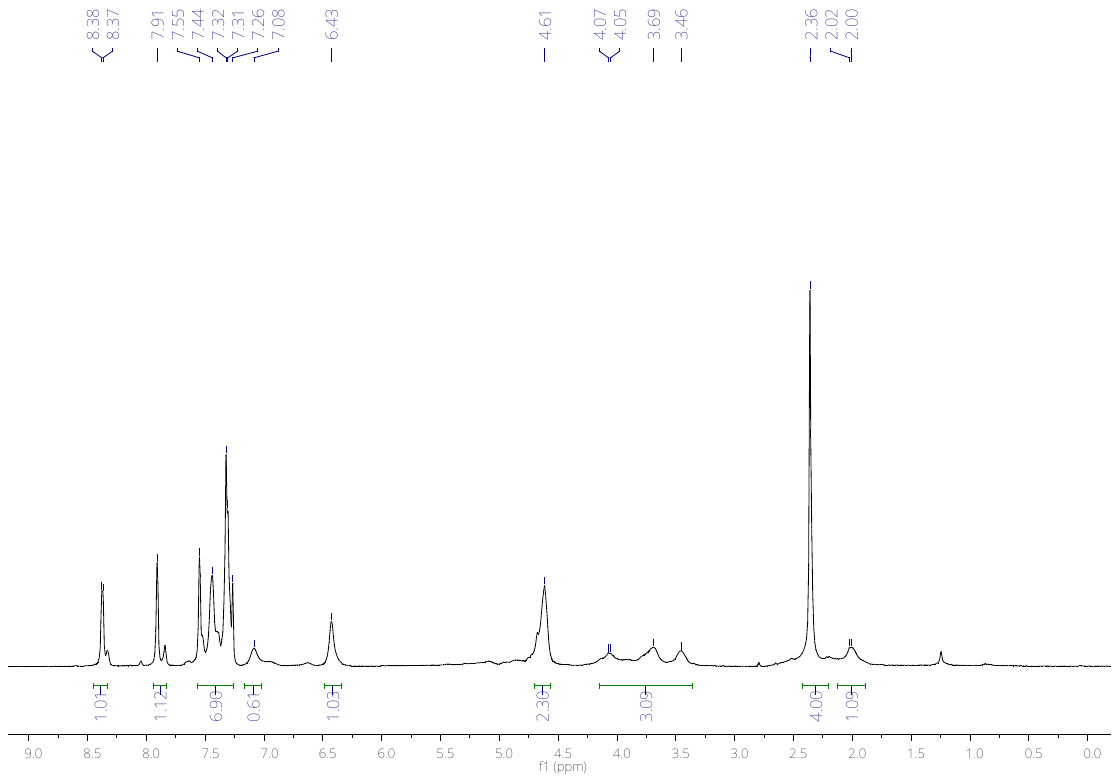

^1^H NMR (CDCl_3_, 400Hz) **10h**

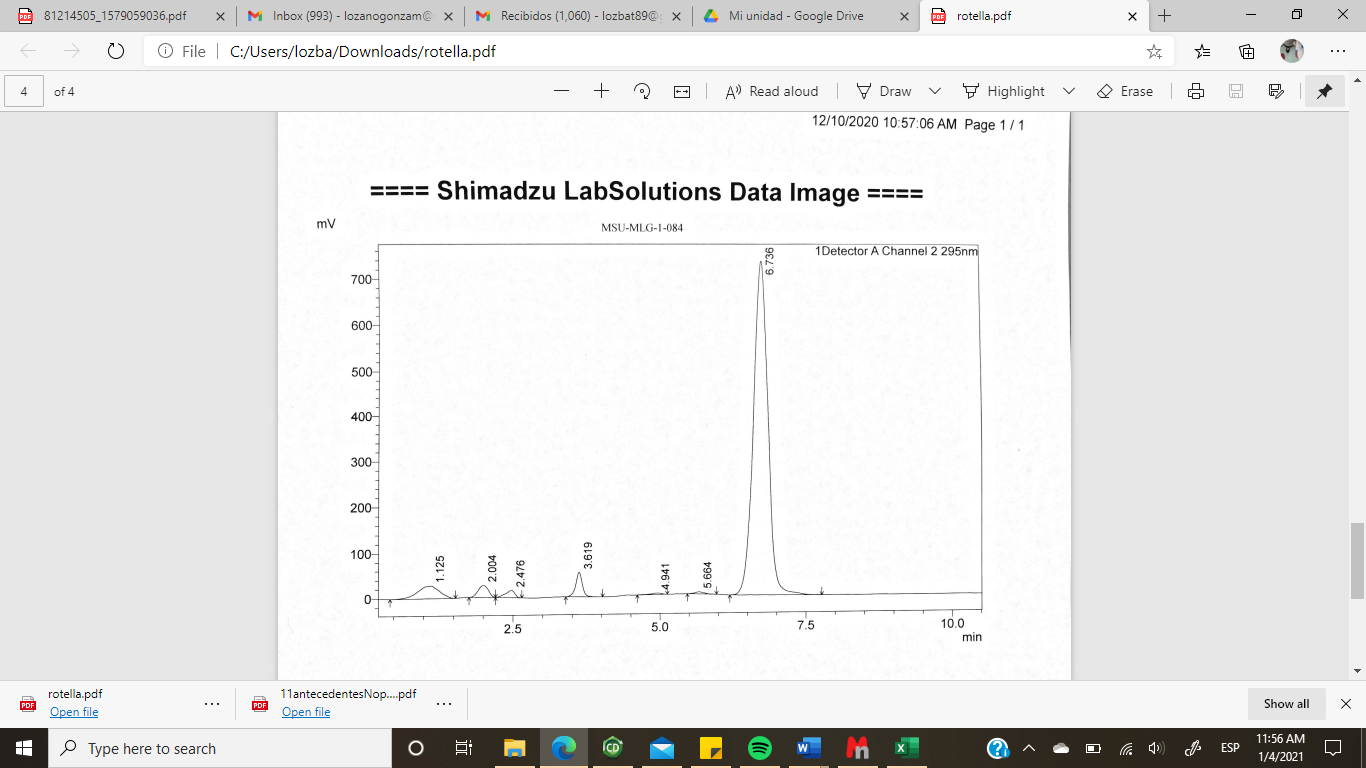

LC Spectrum **10h**

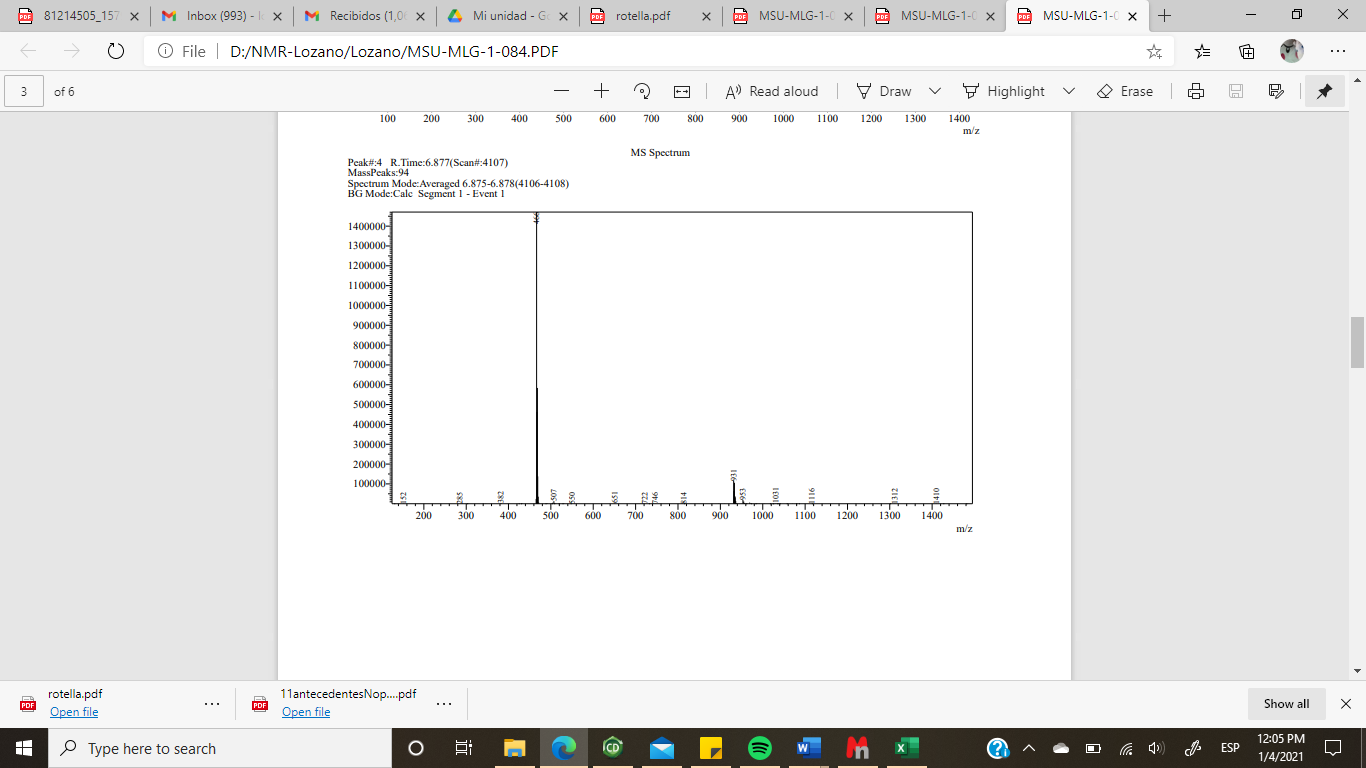

**MS Spectrum** **10h** (M+ H)^+^ +(MeOH + H)^+^ 466 m/z

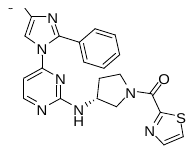

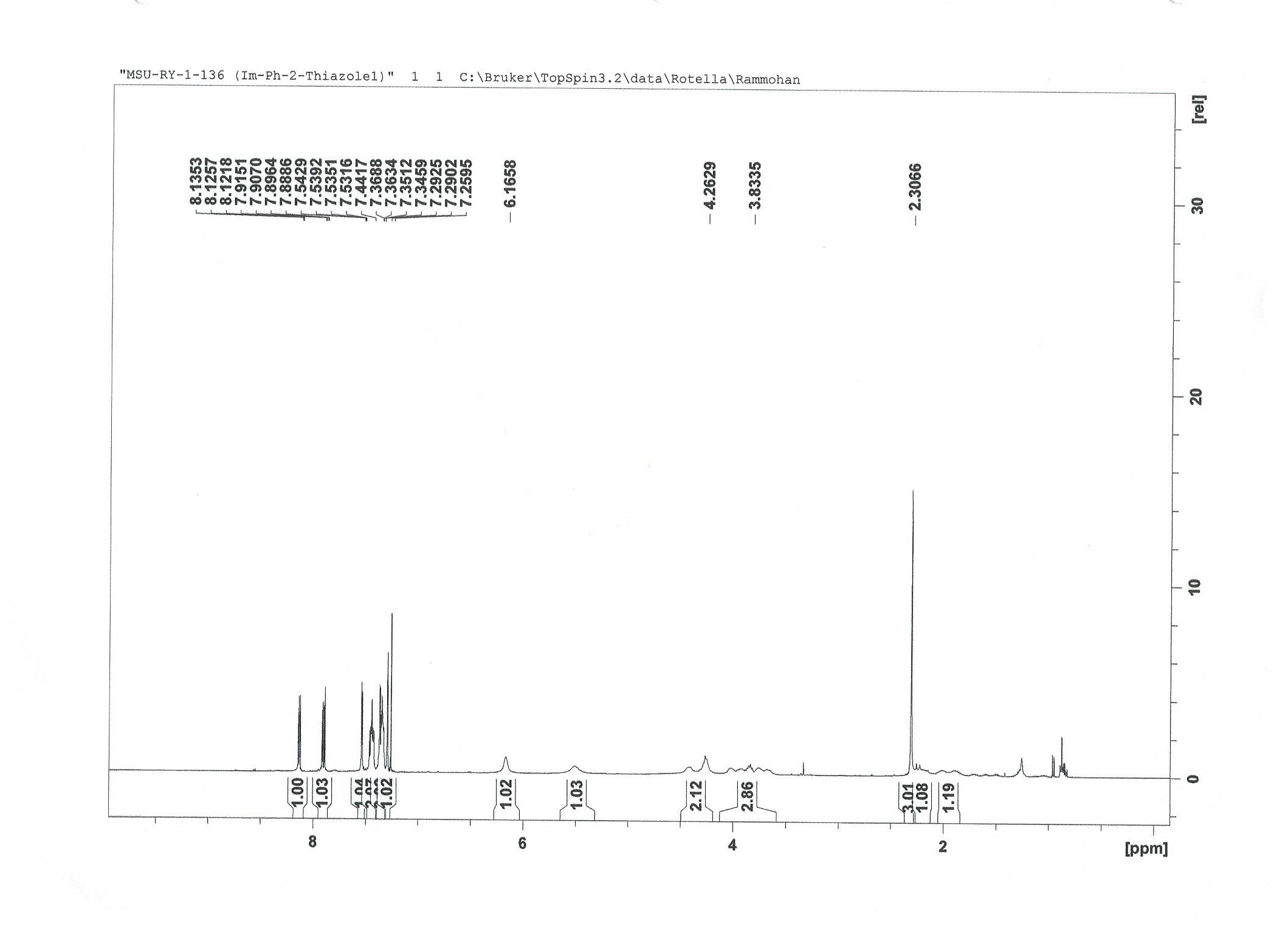

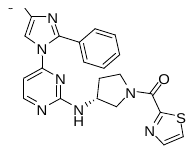

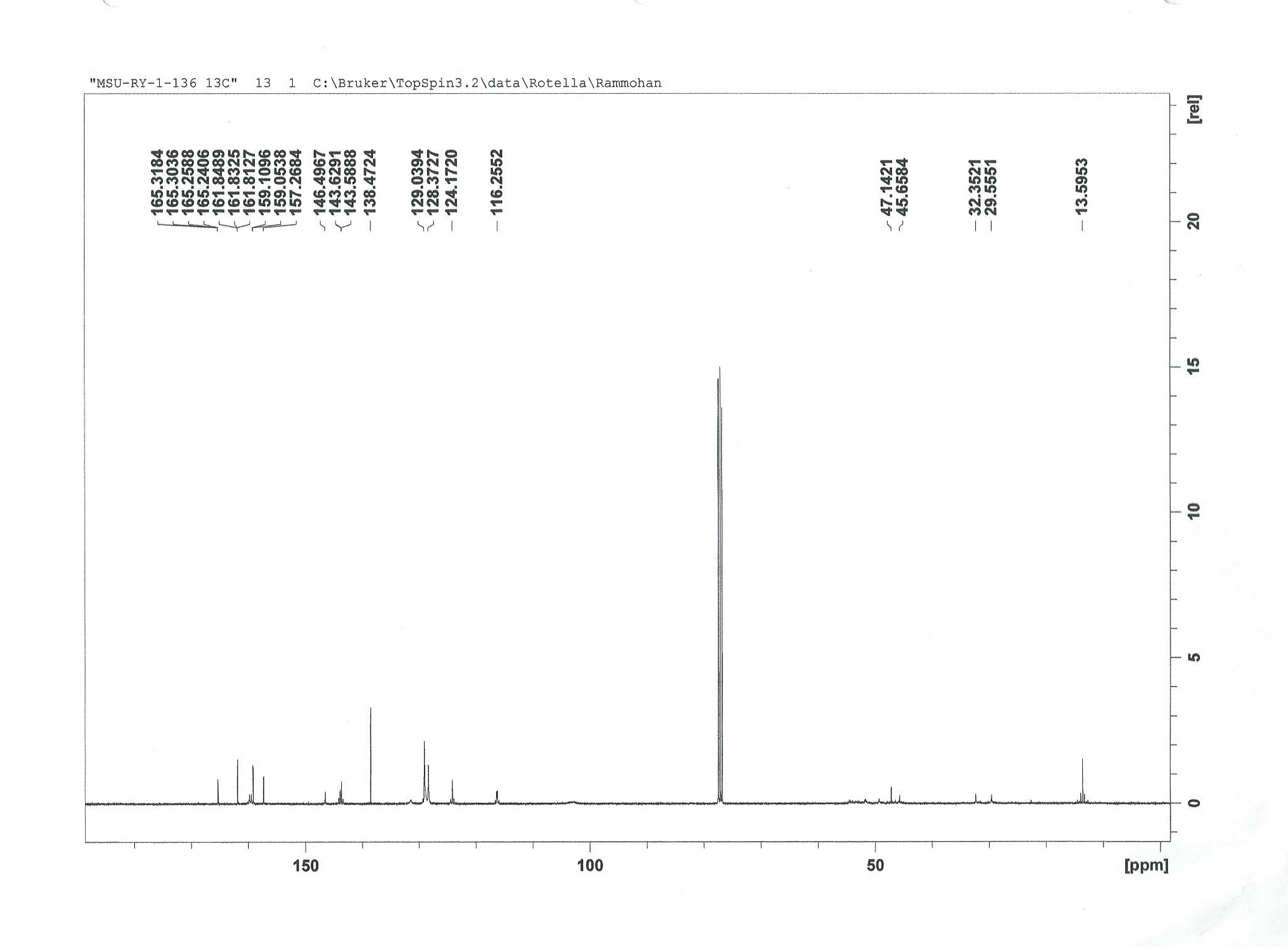

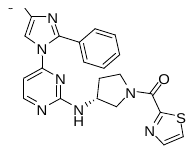

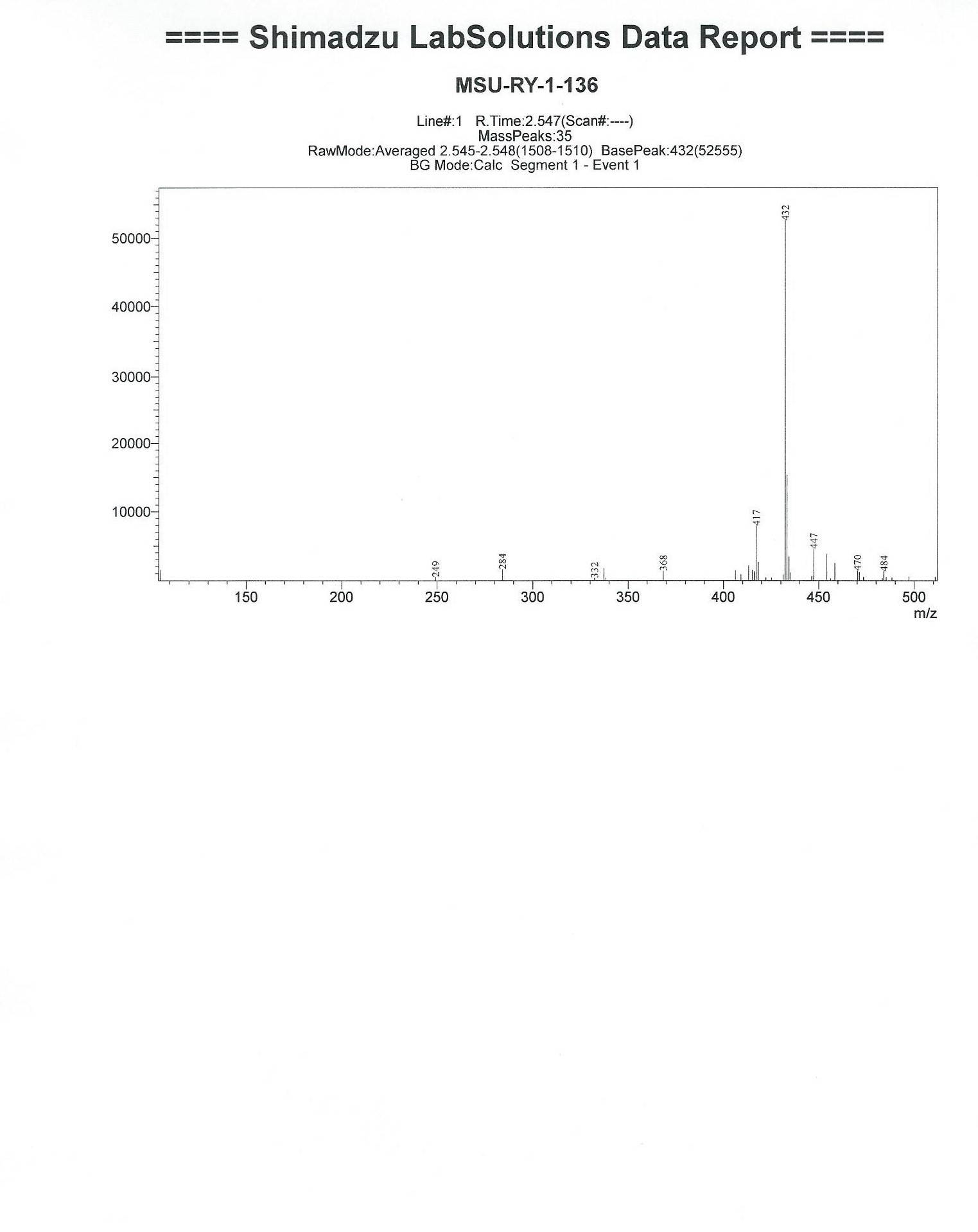

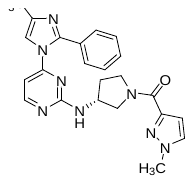

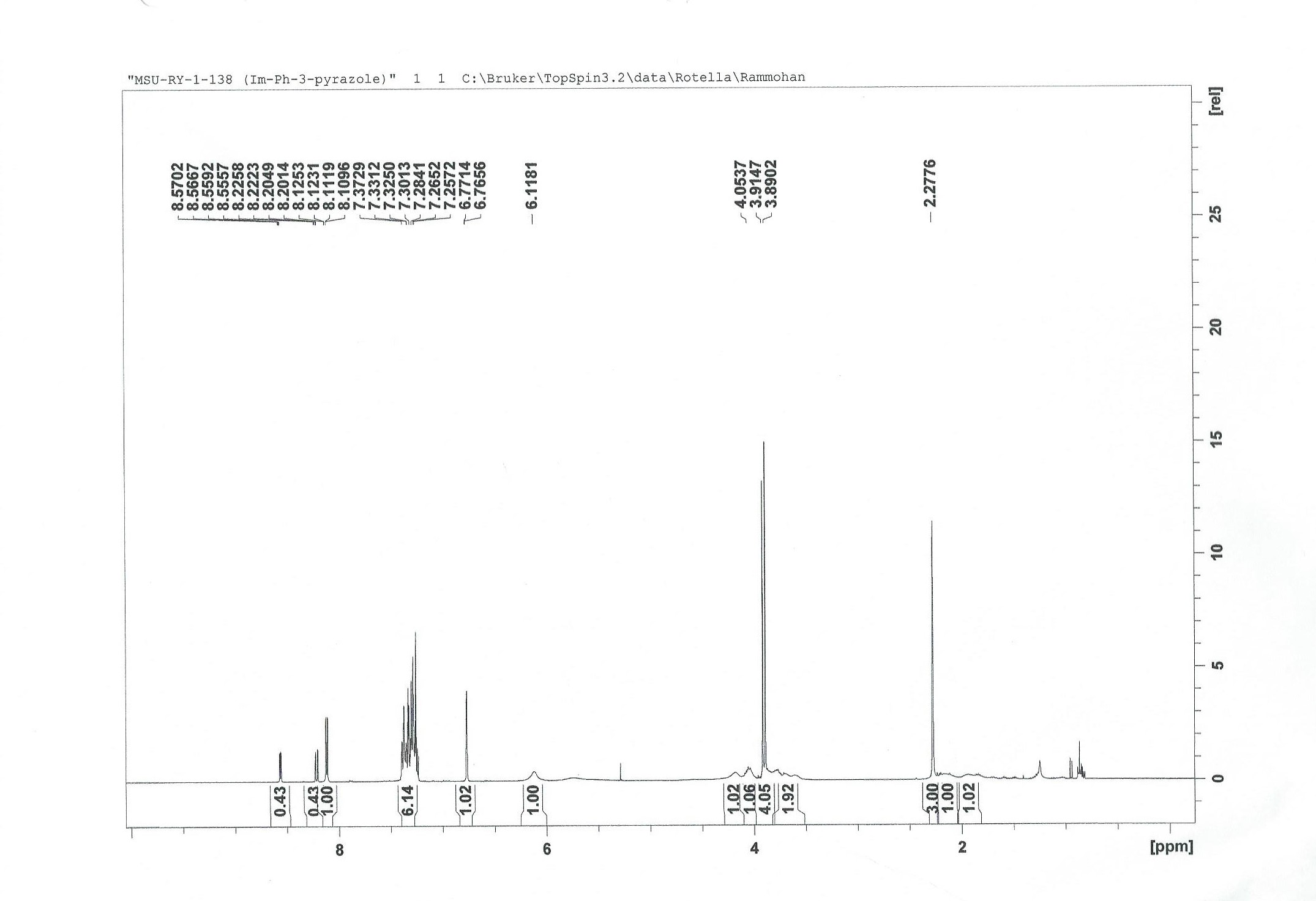

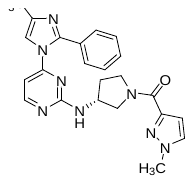

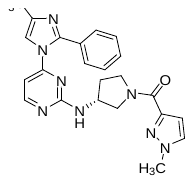

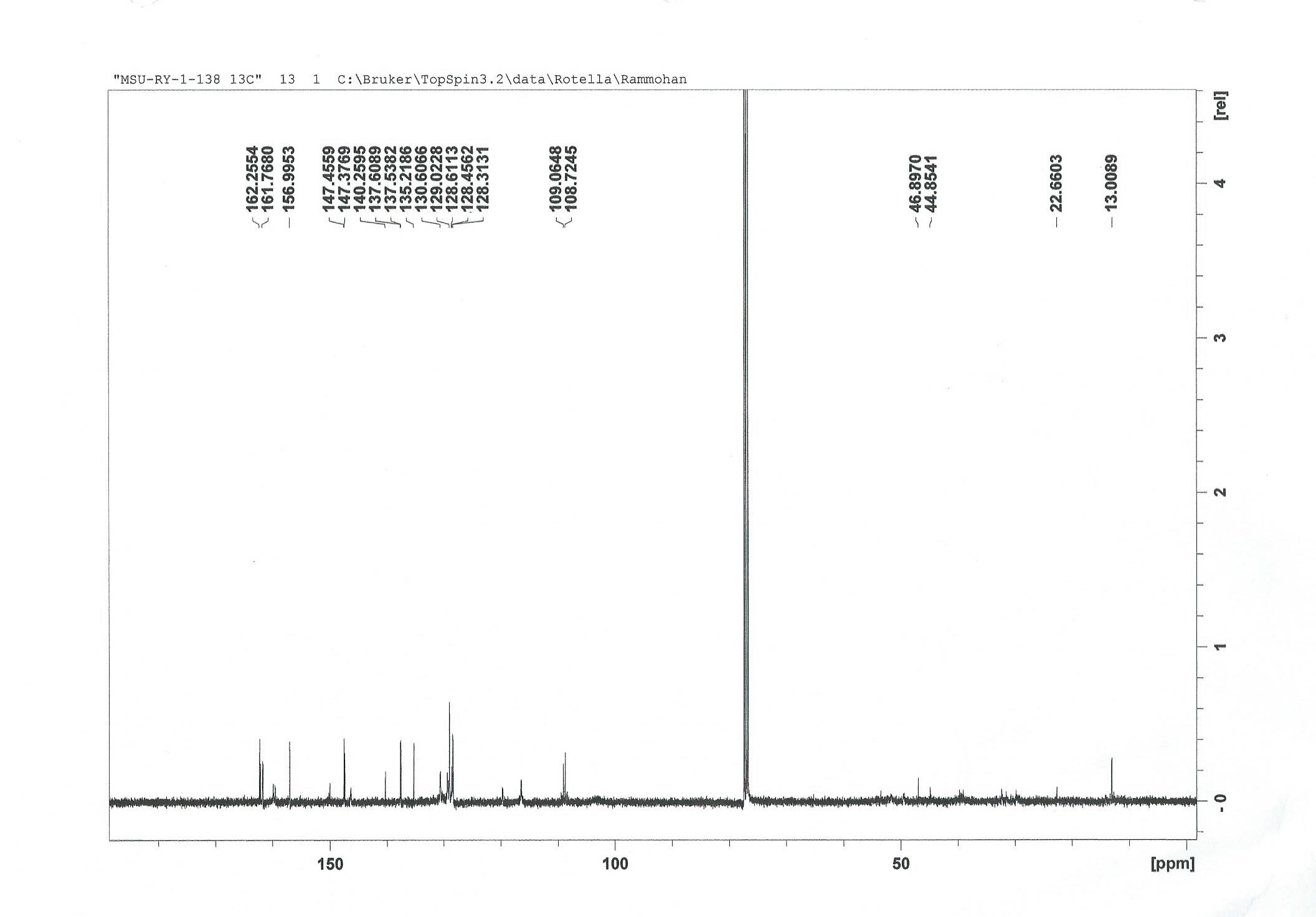

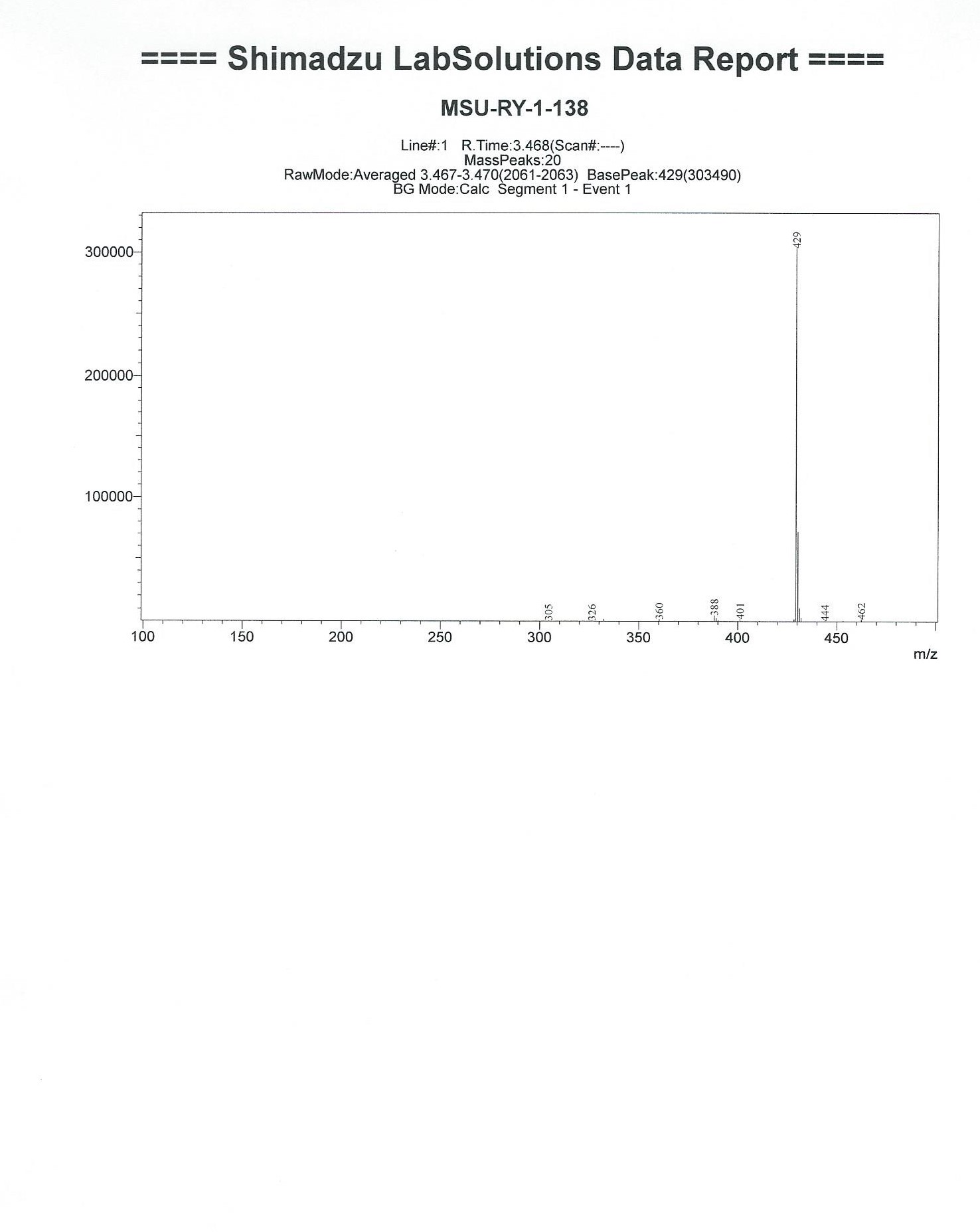

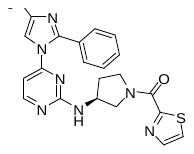

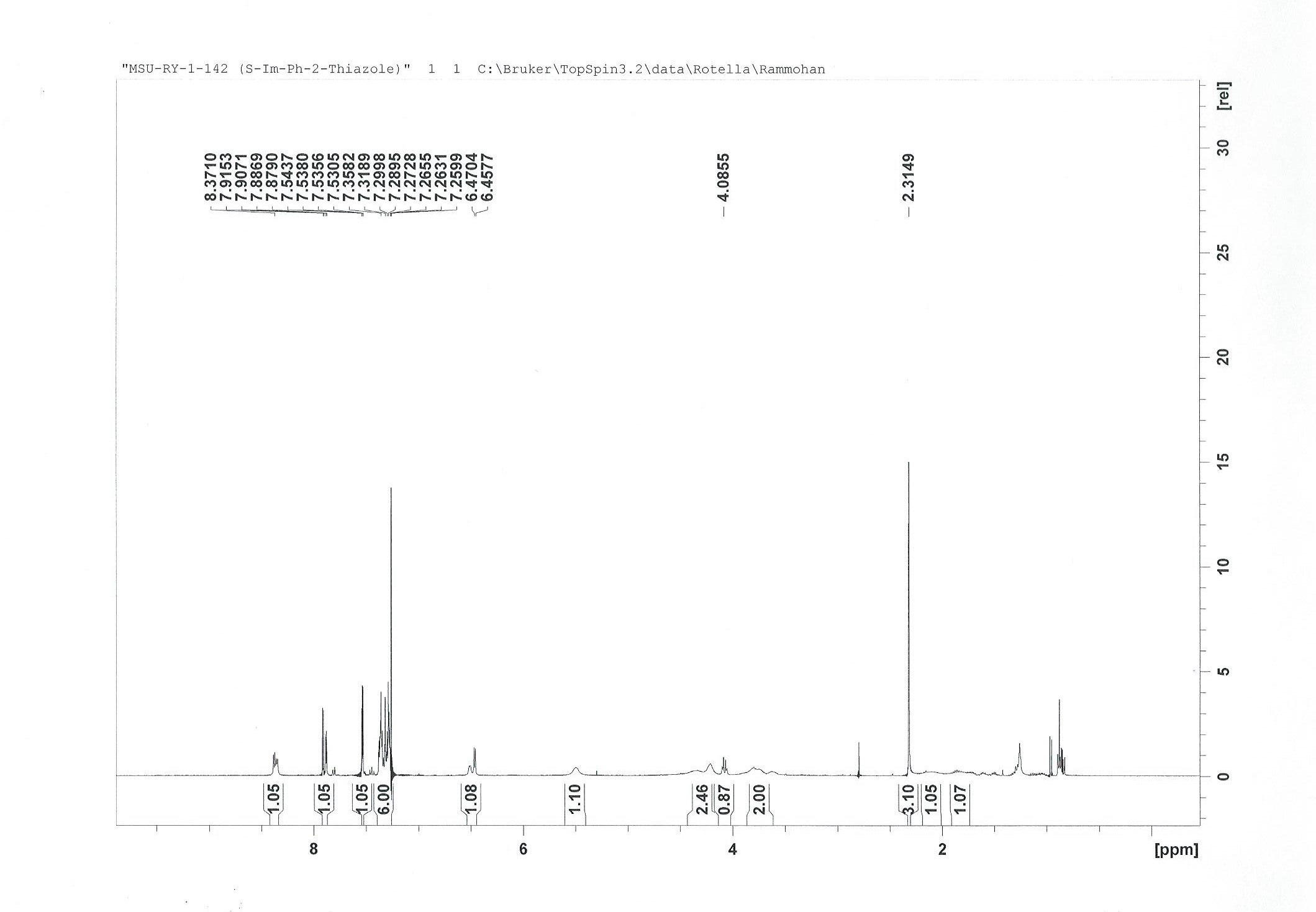

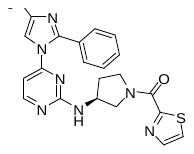

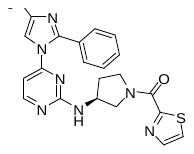

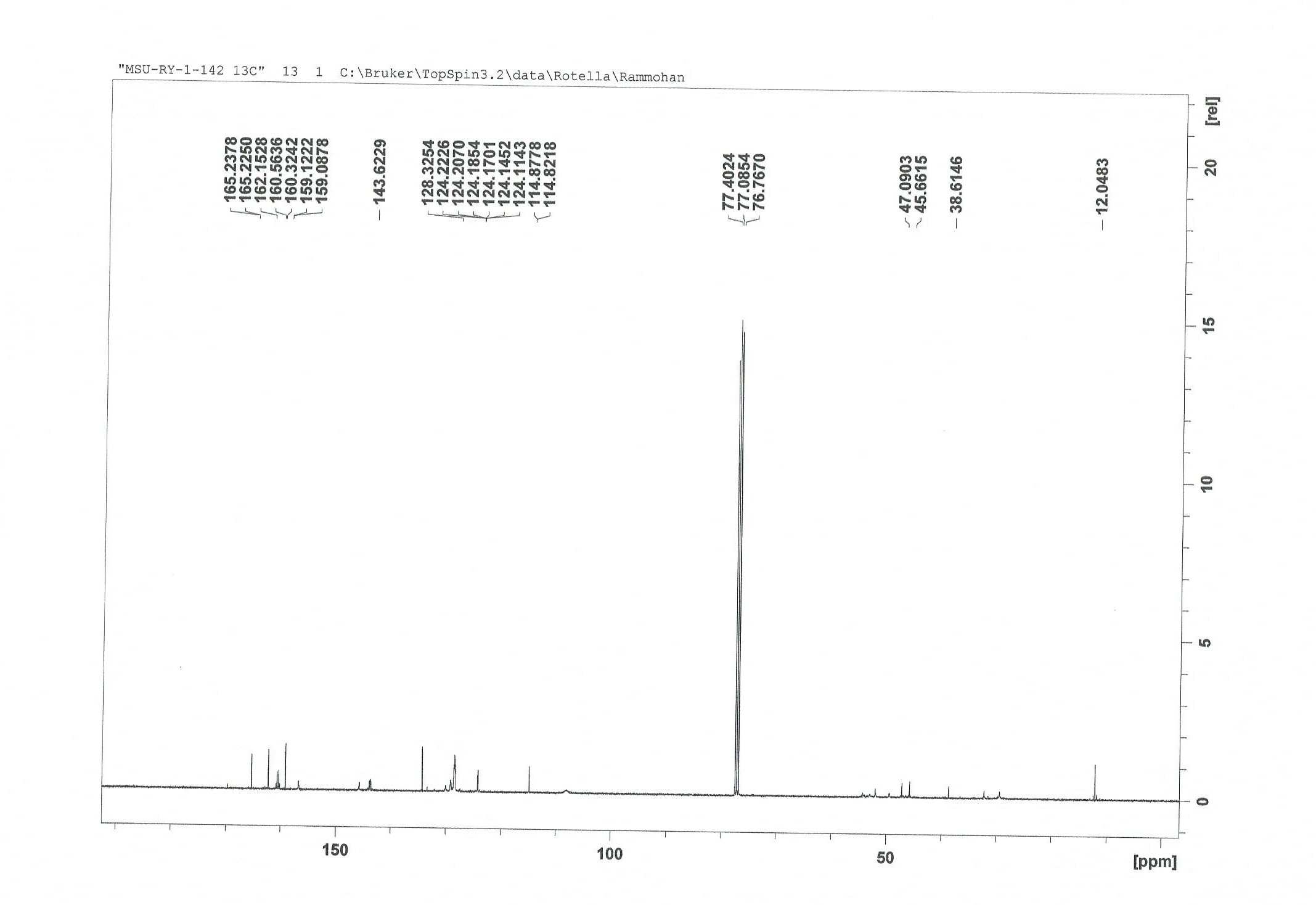

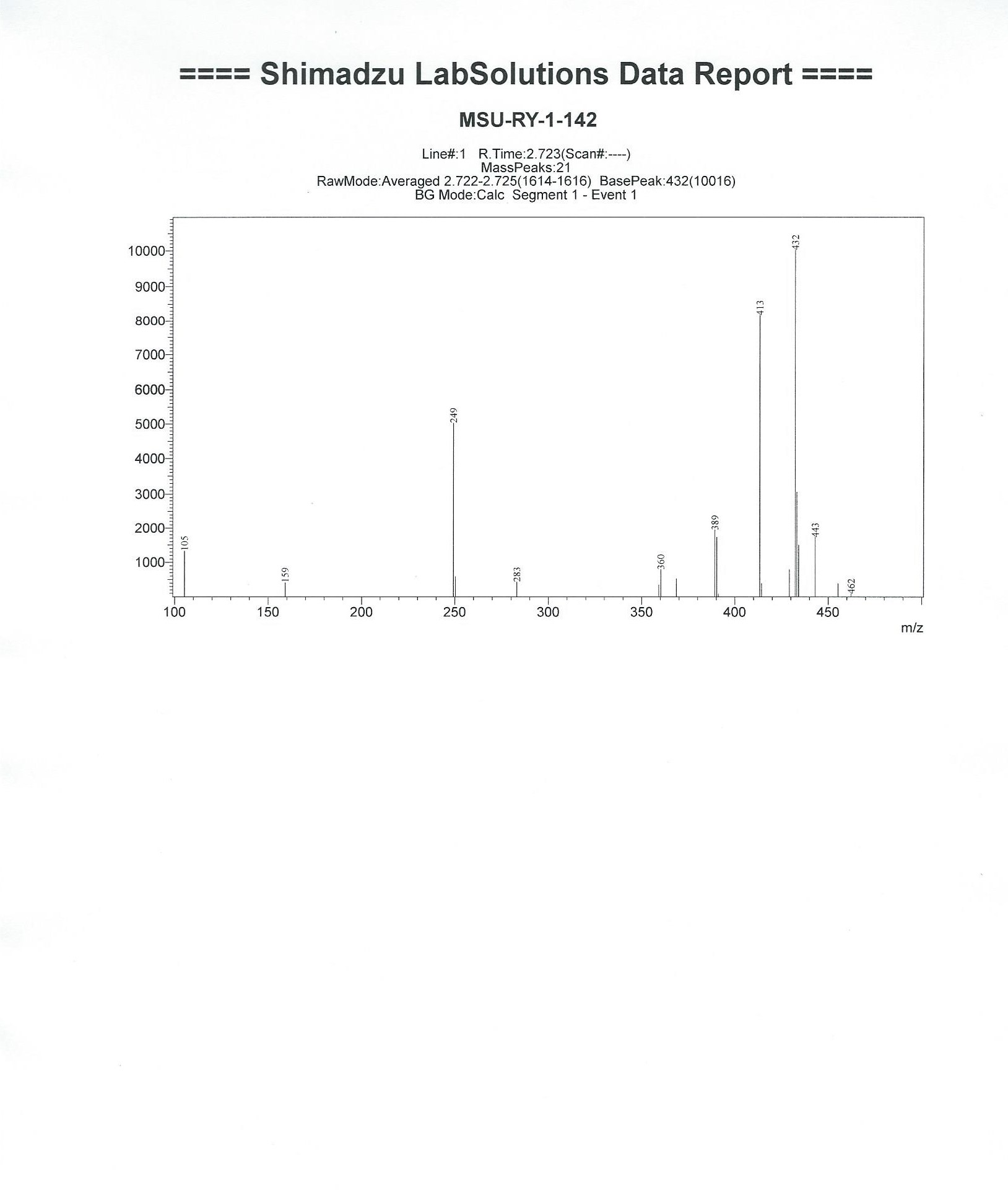

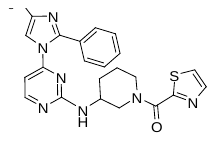

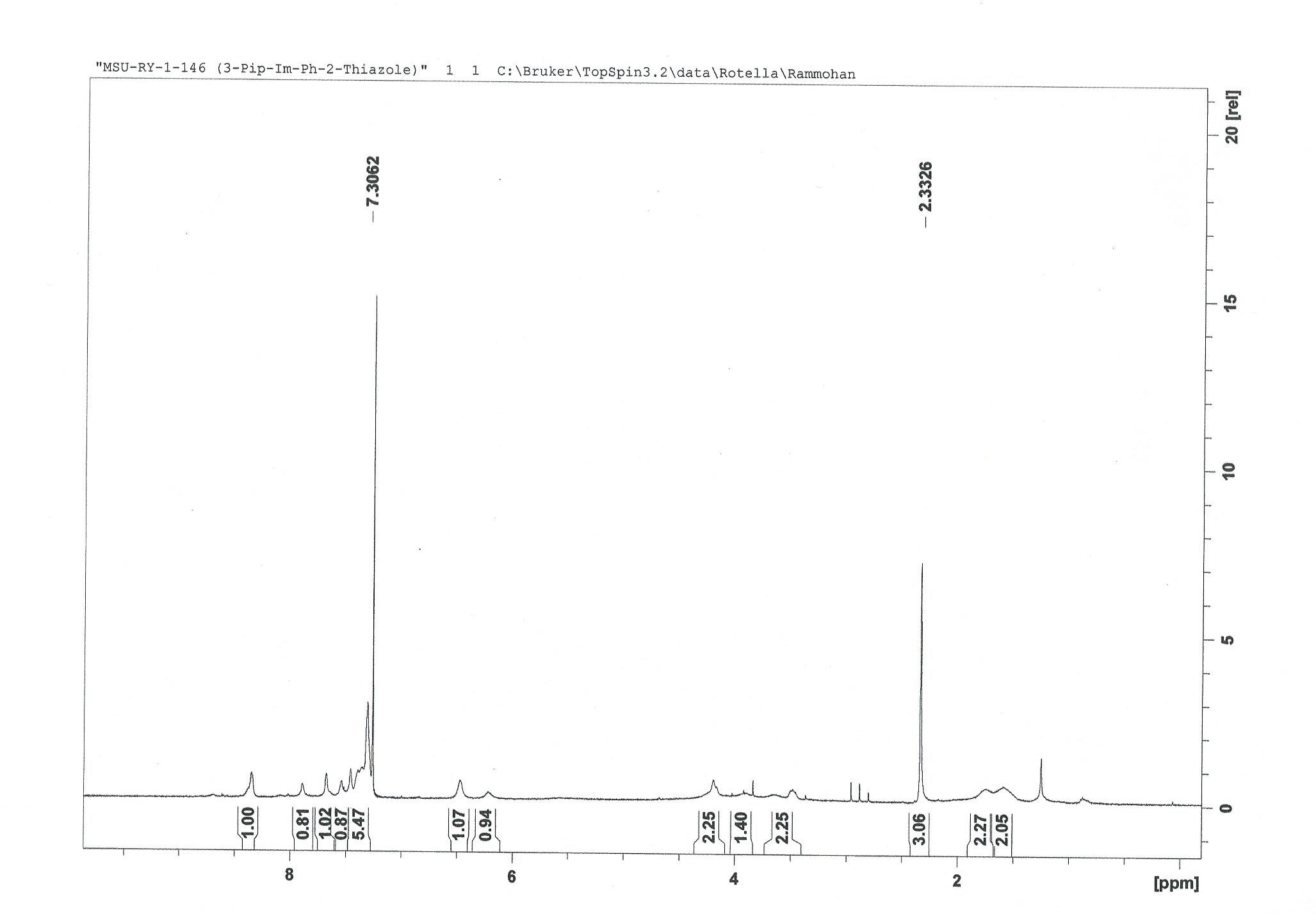

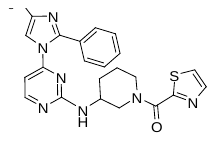

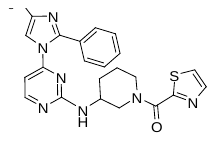

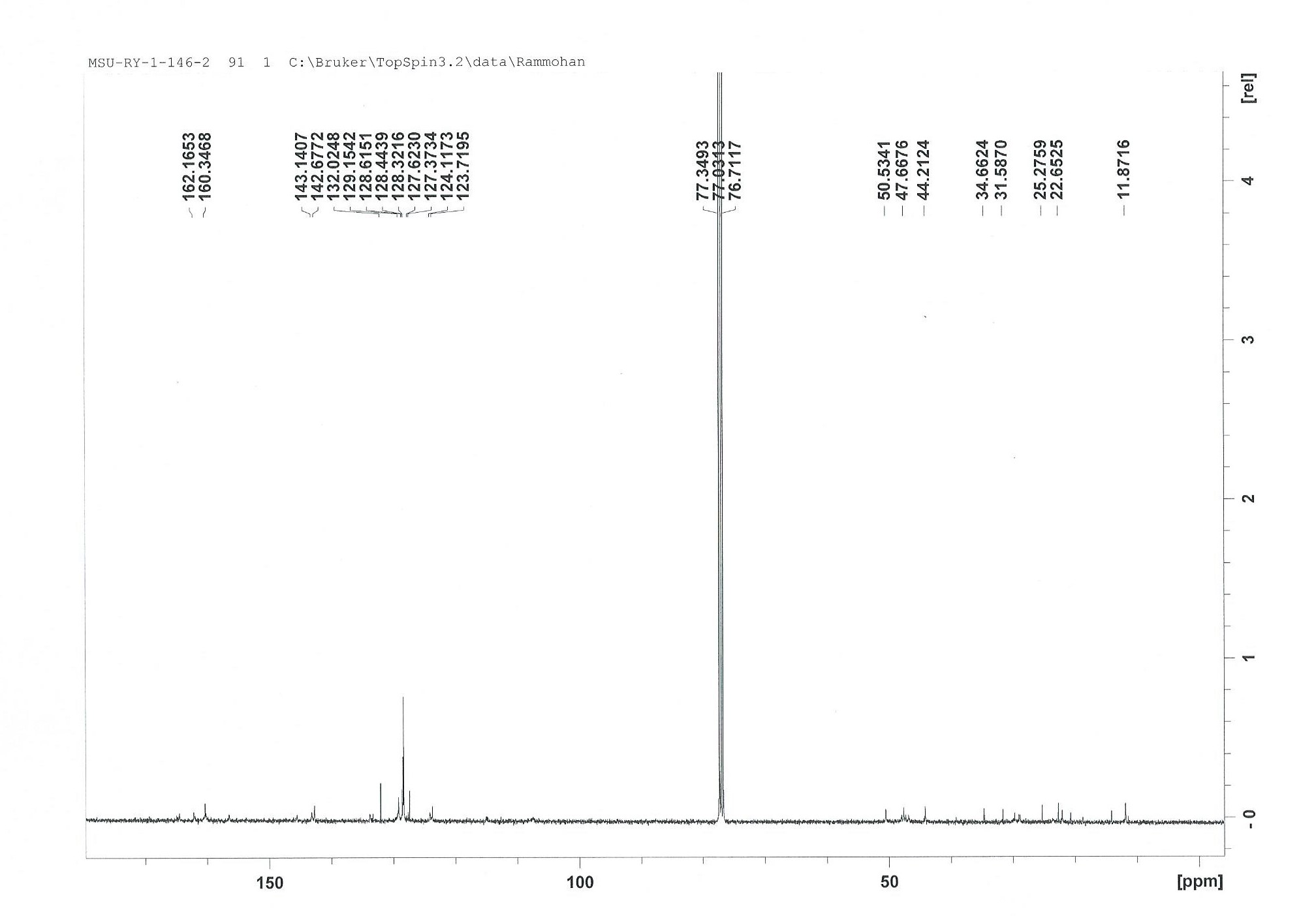

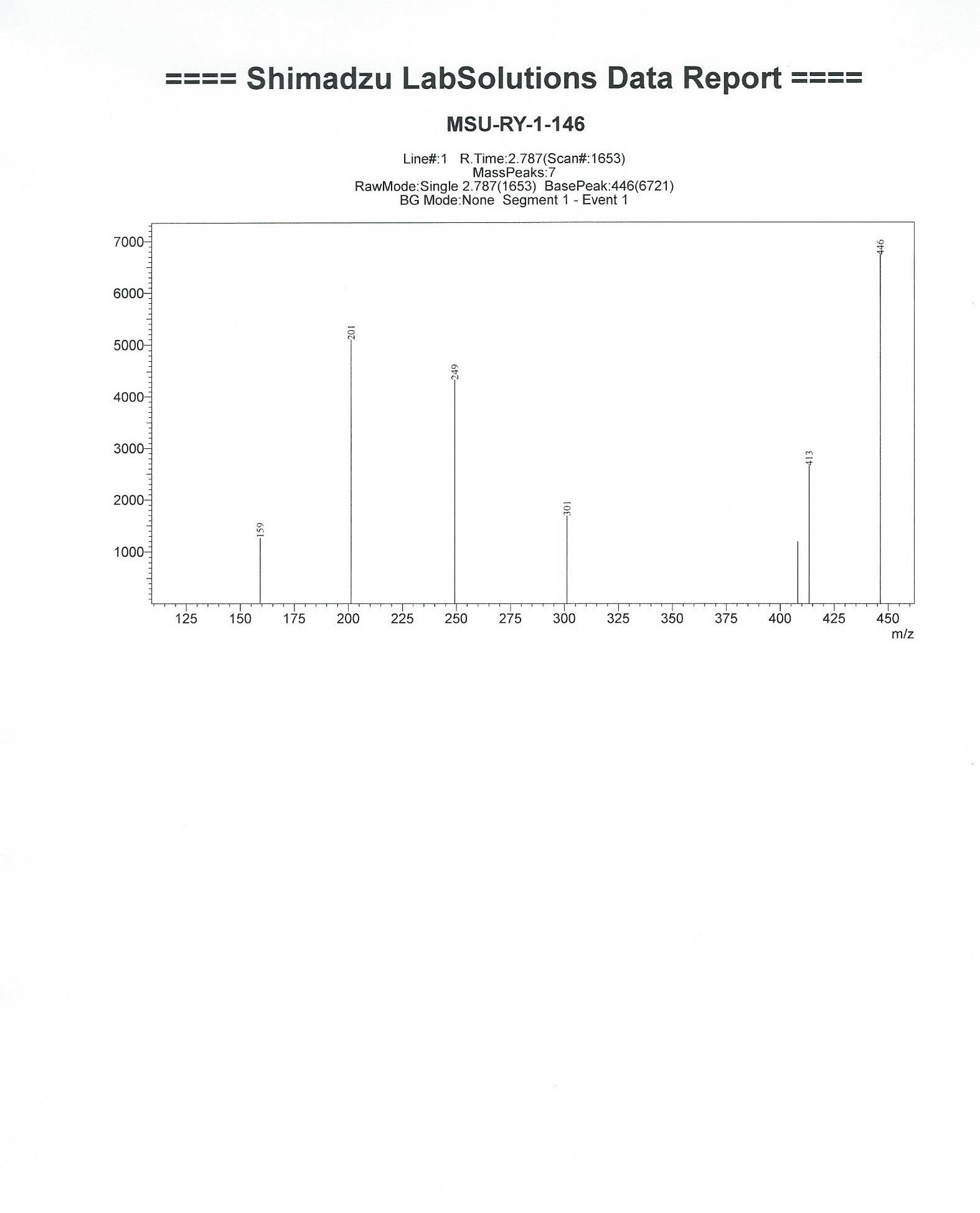

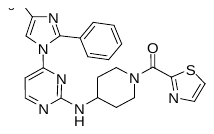

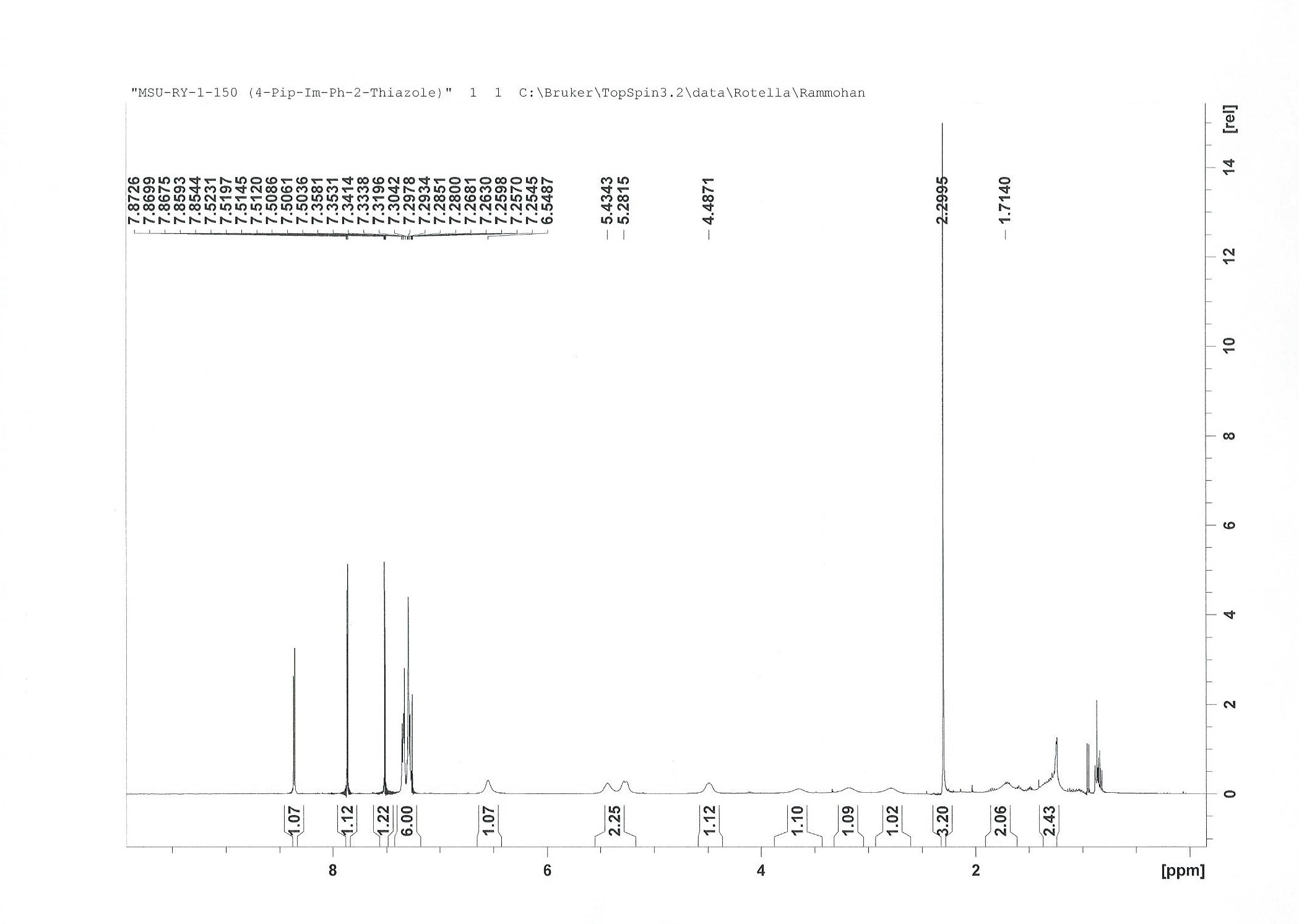

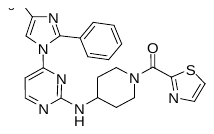

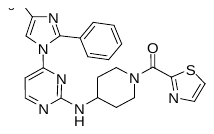

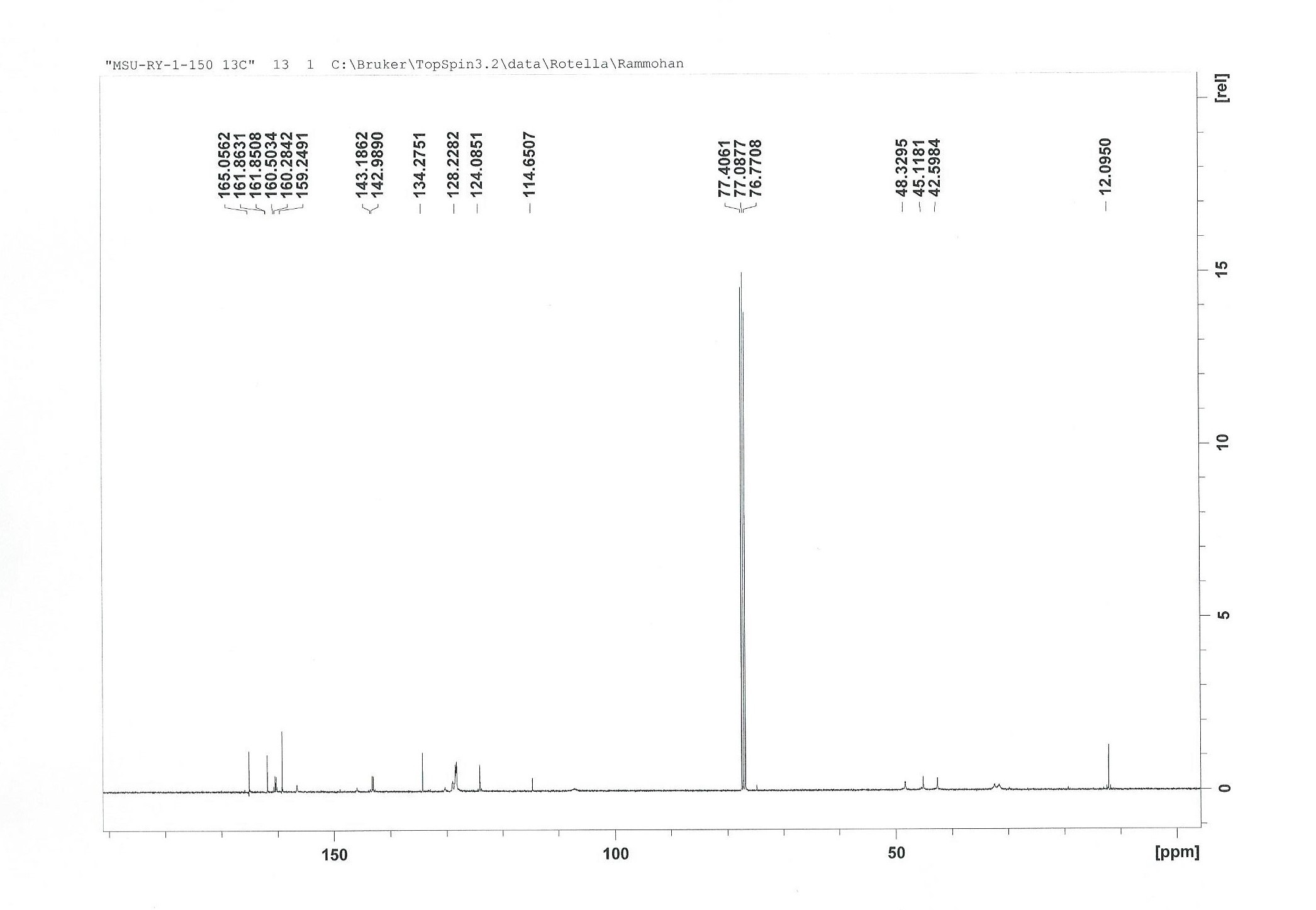

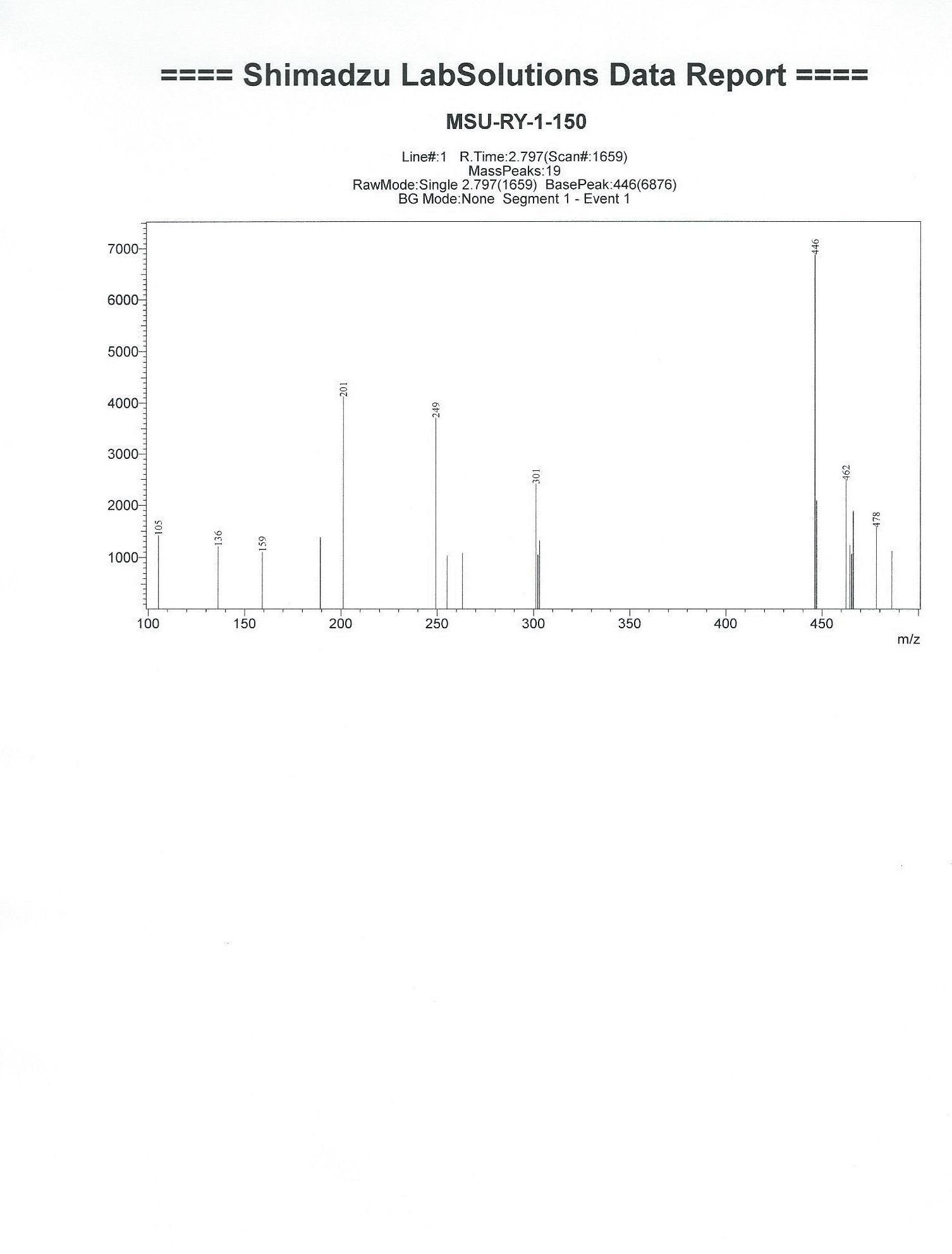

**S1.6.** ^1^H, ^13^C NMR and Mass spectra of **MSU-RY-1-165**

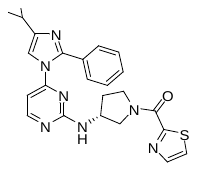

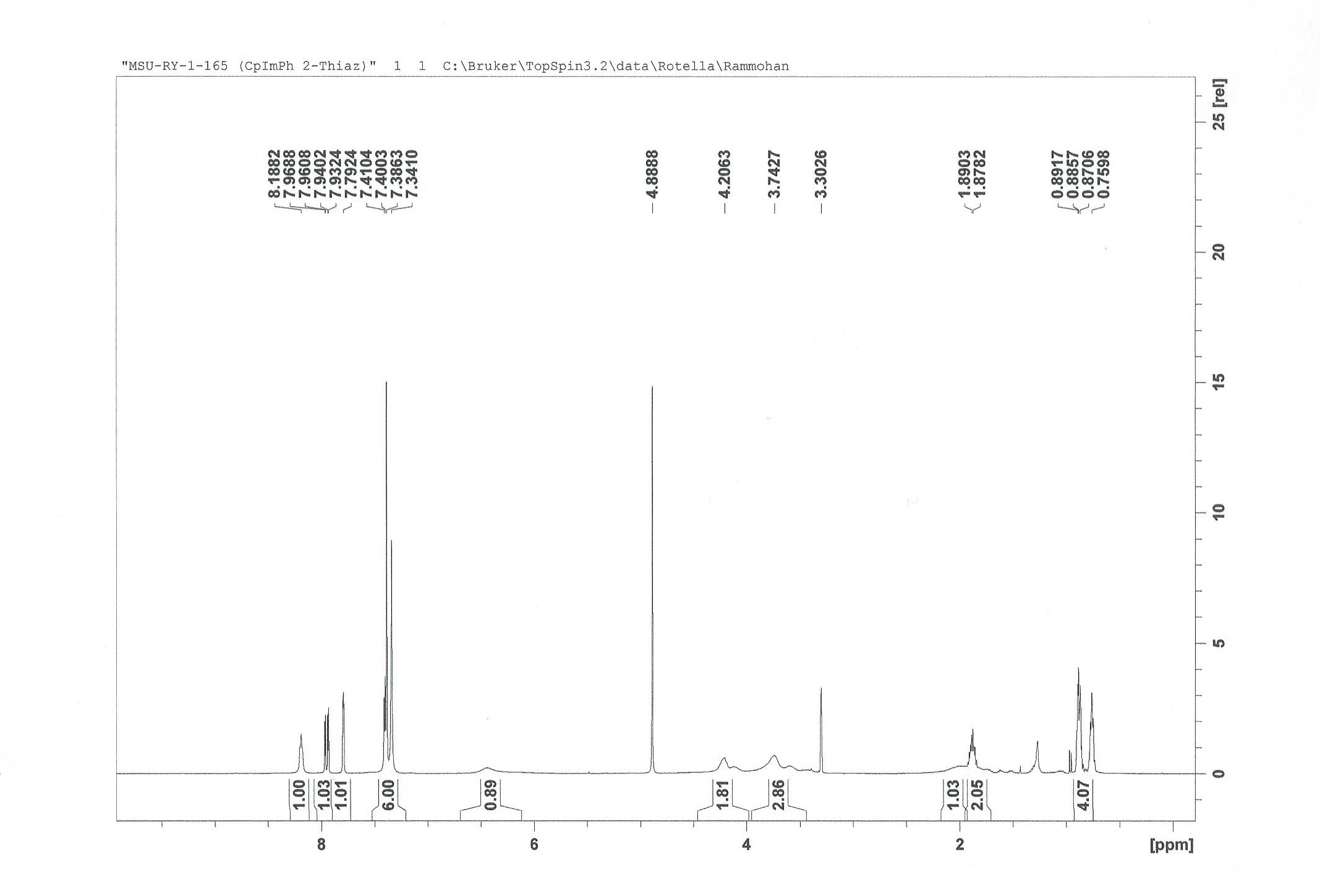

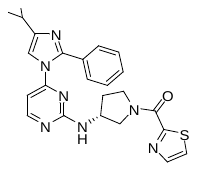

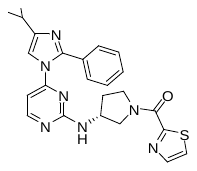

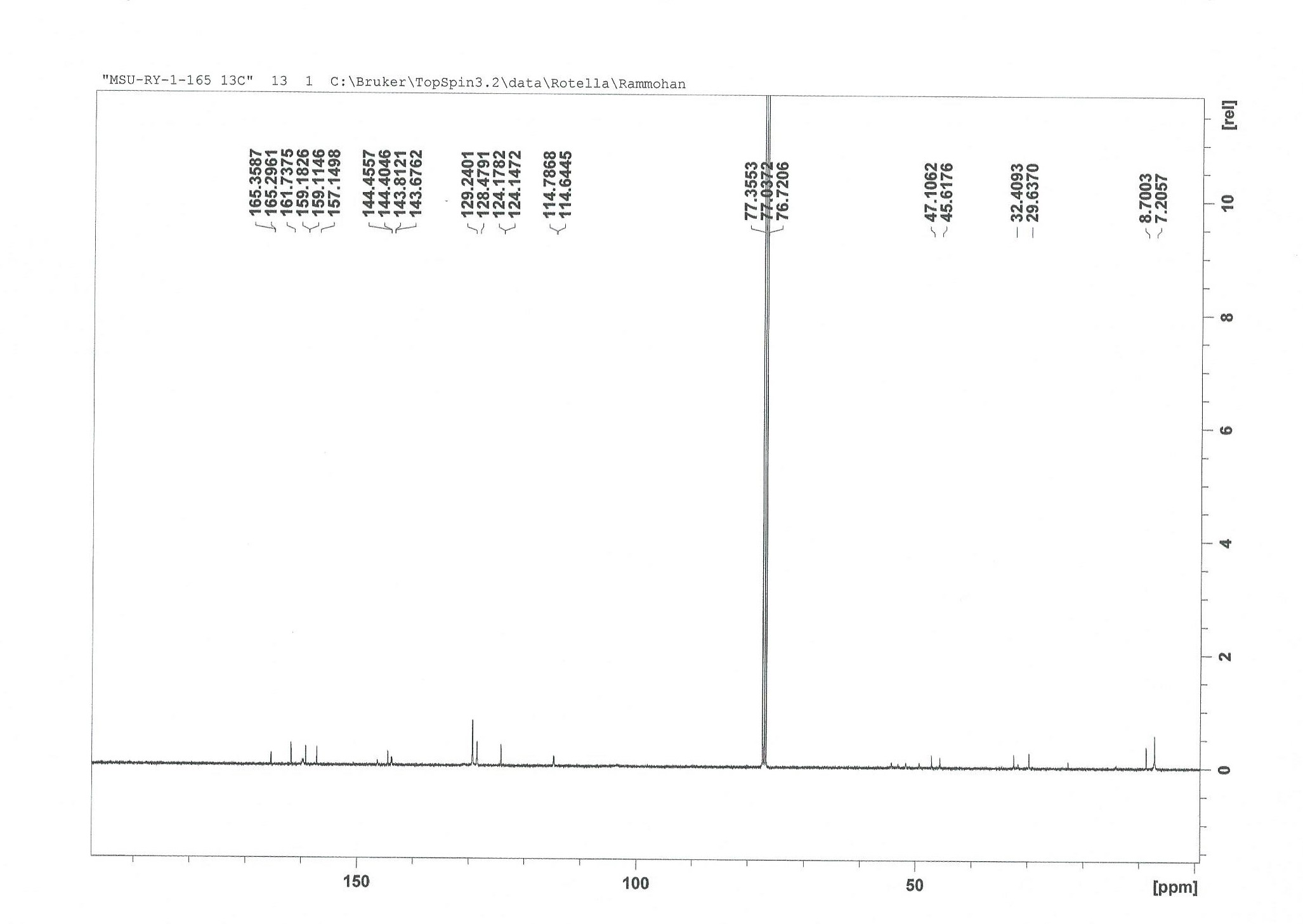

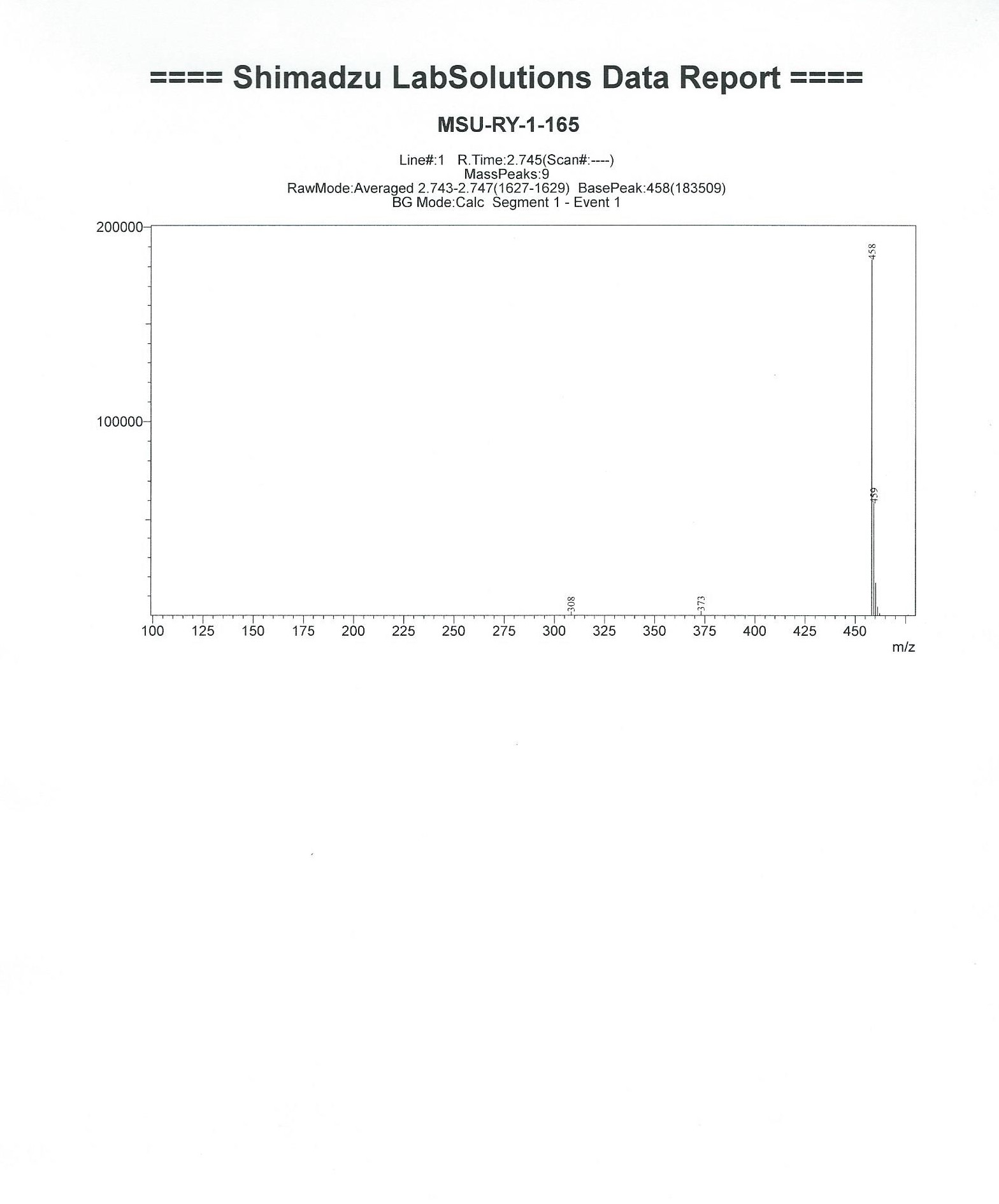

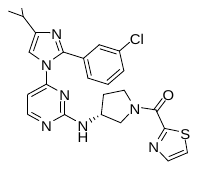

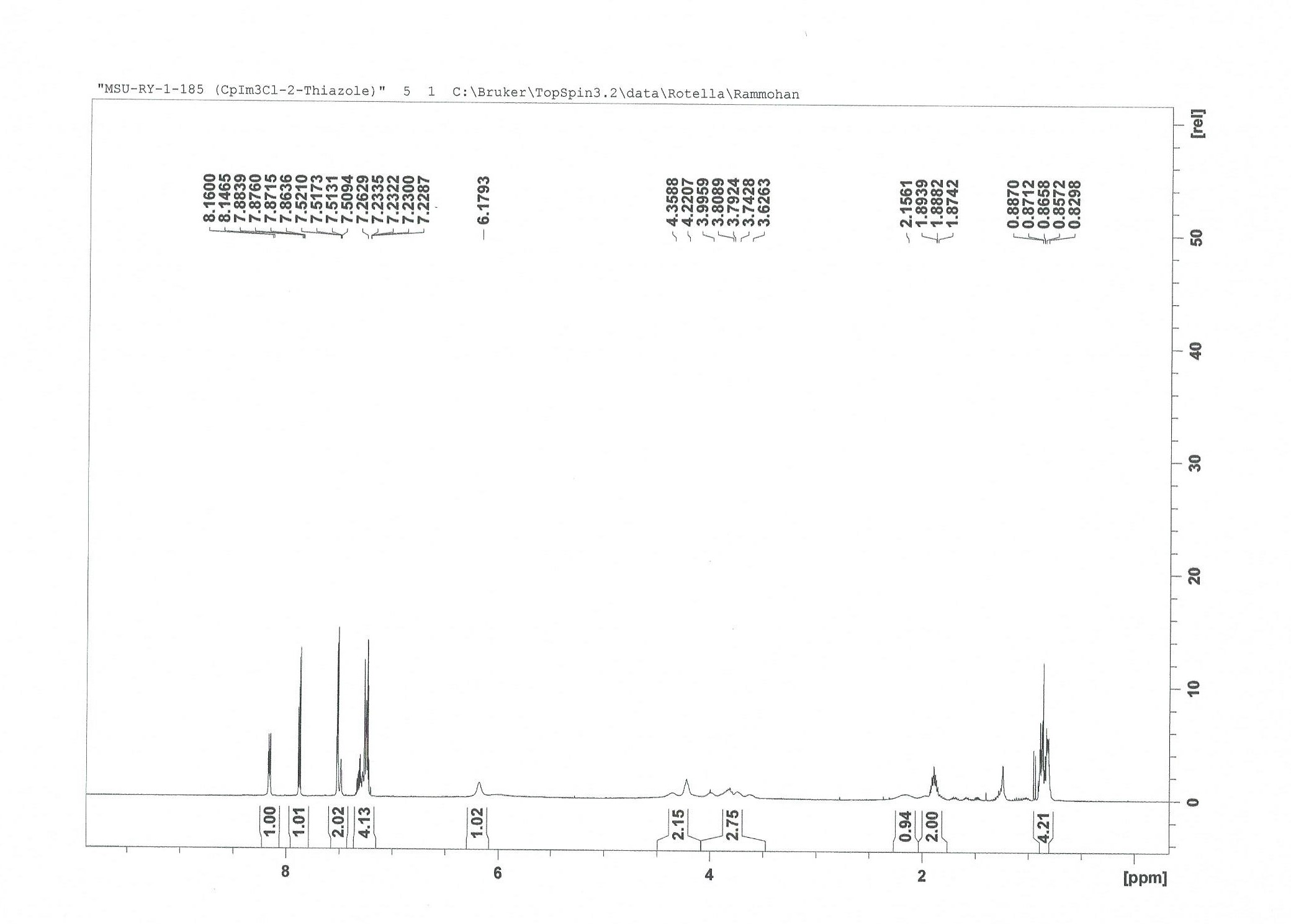

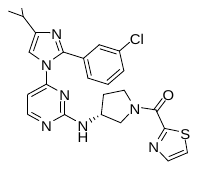

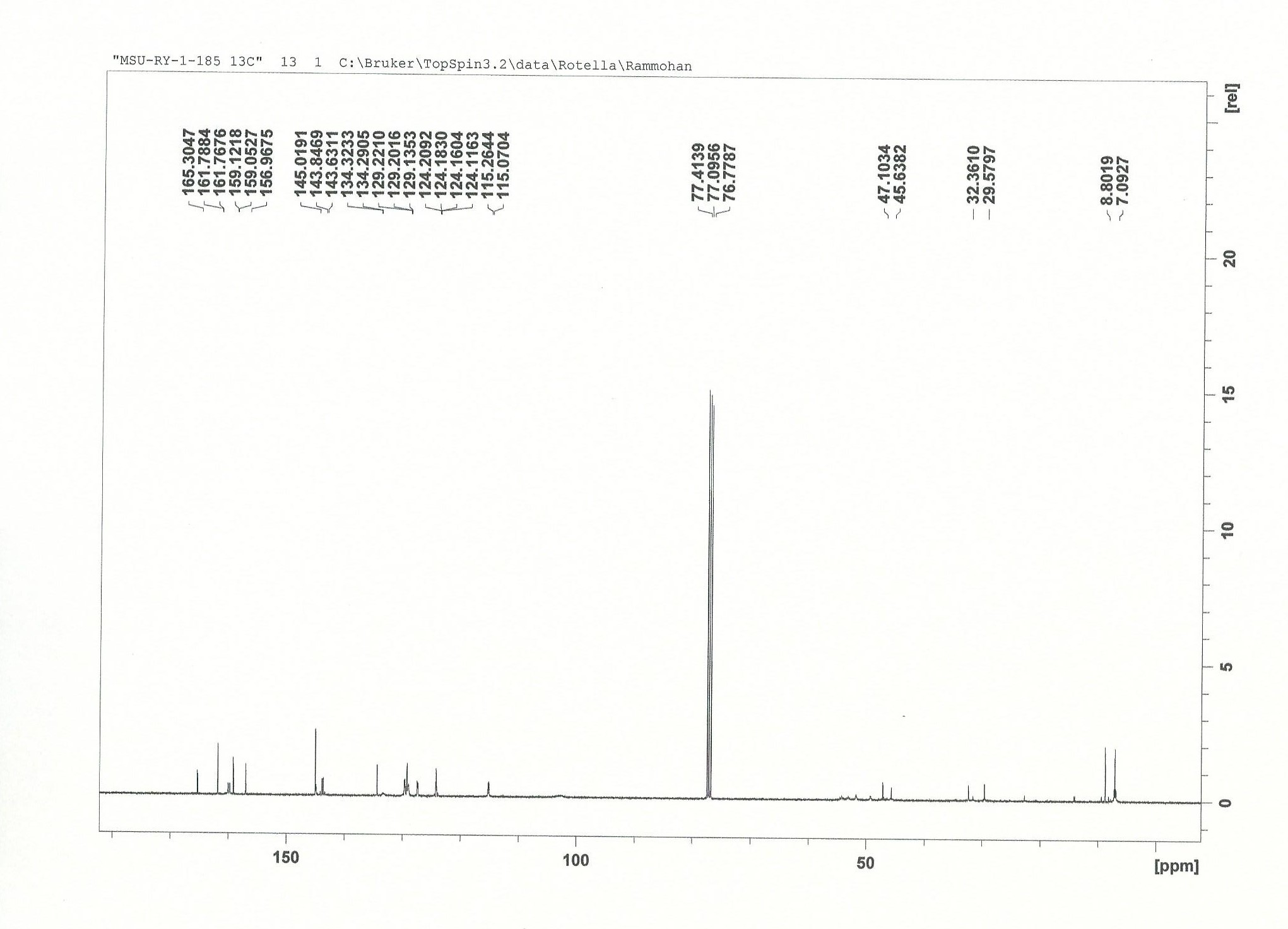

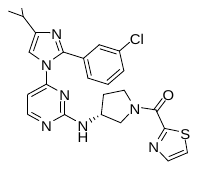

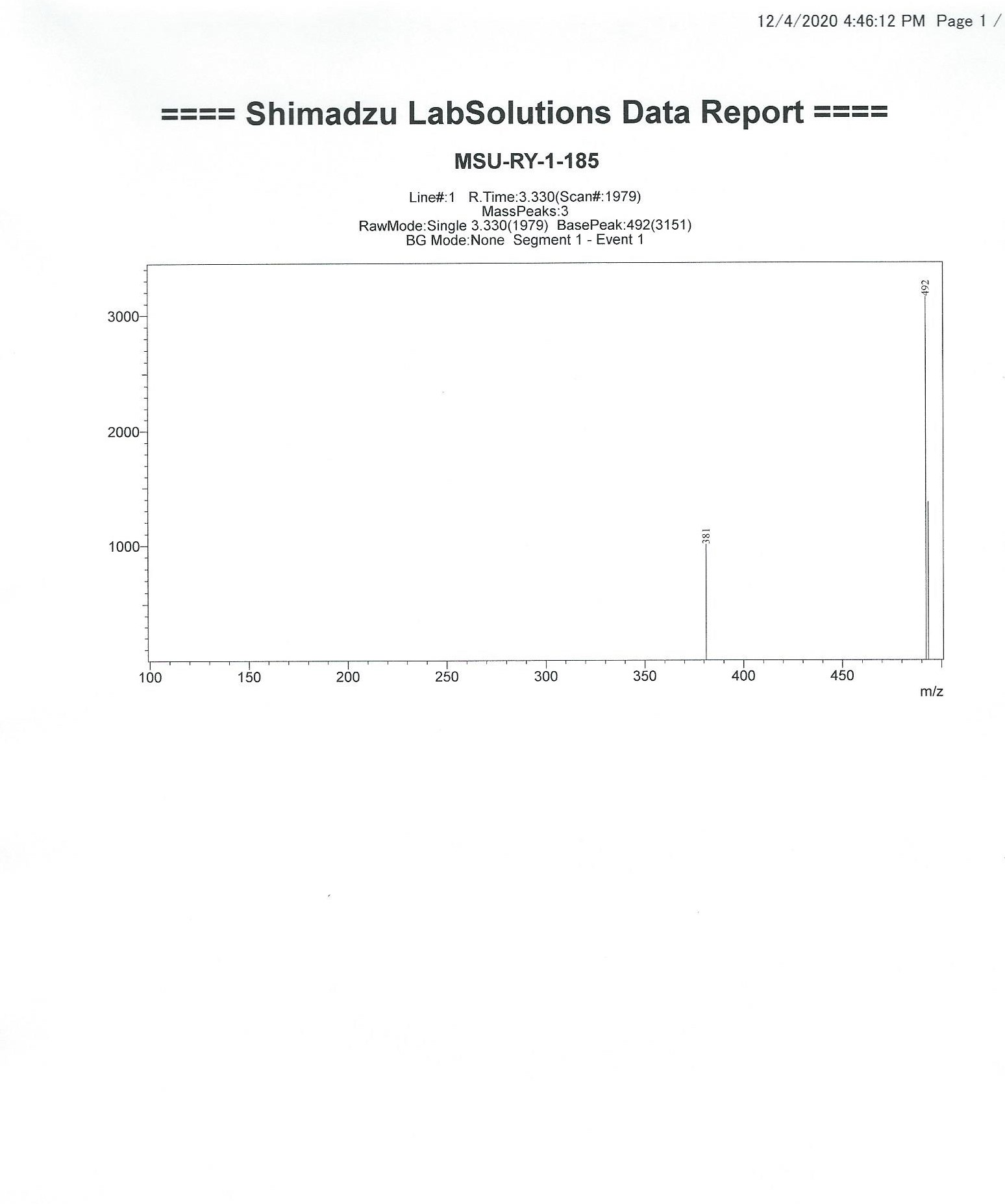

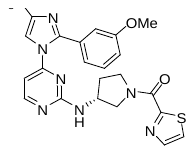

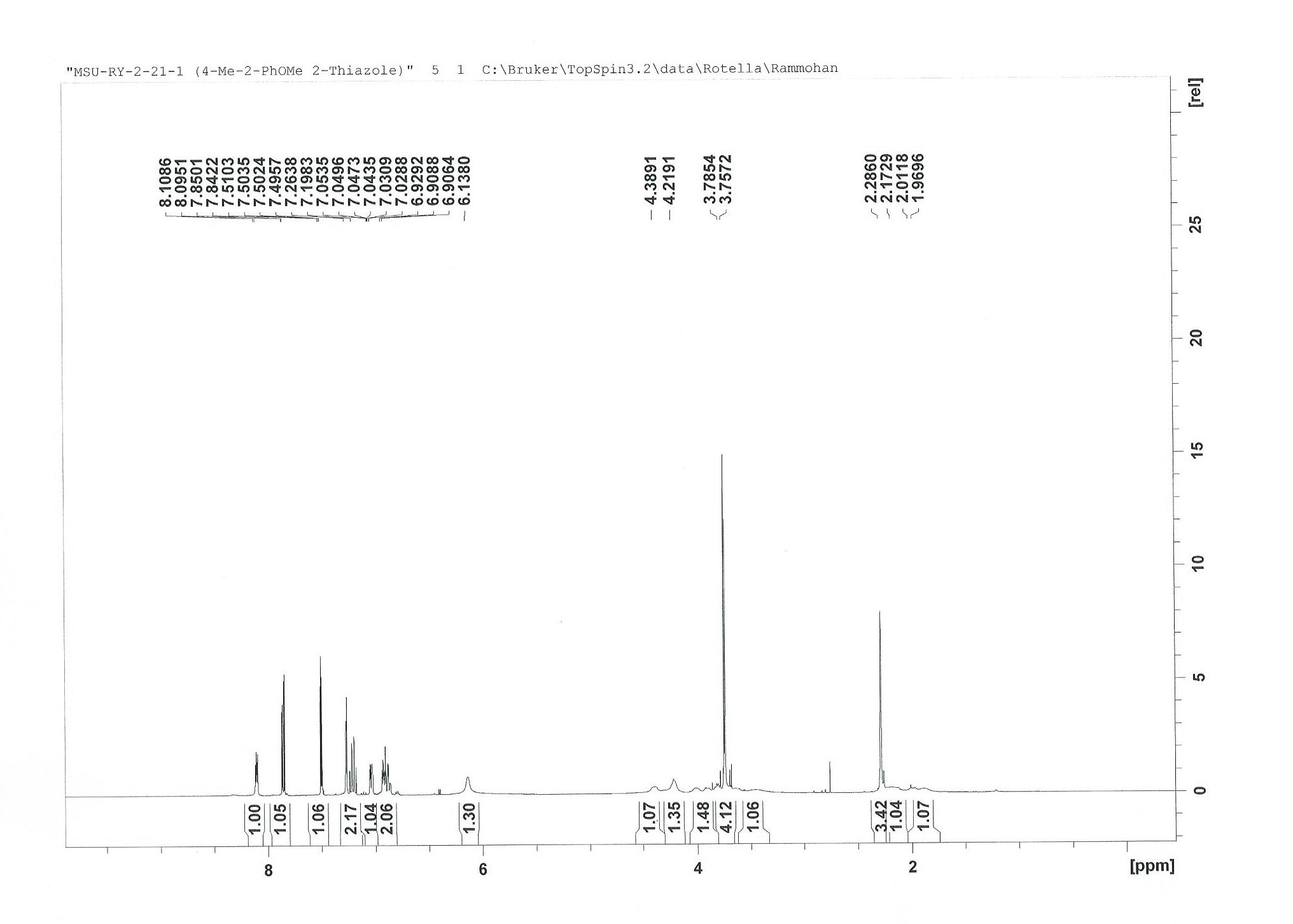

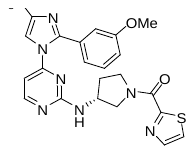

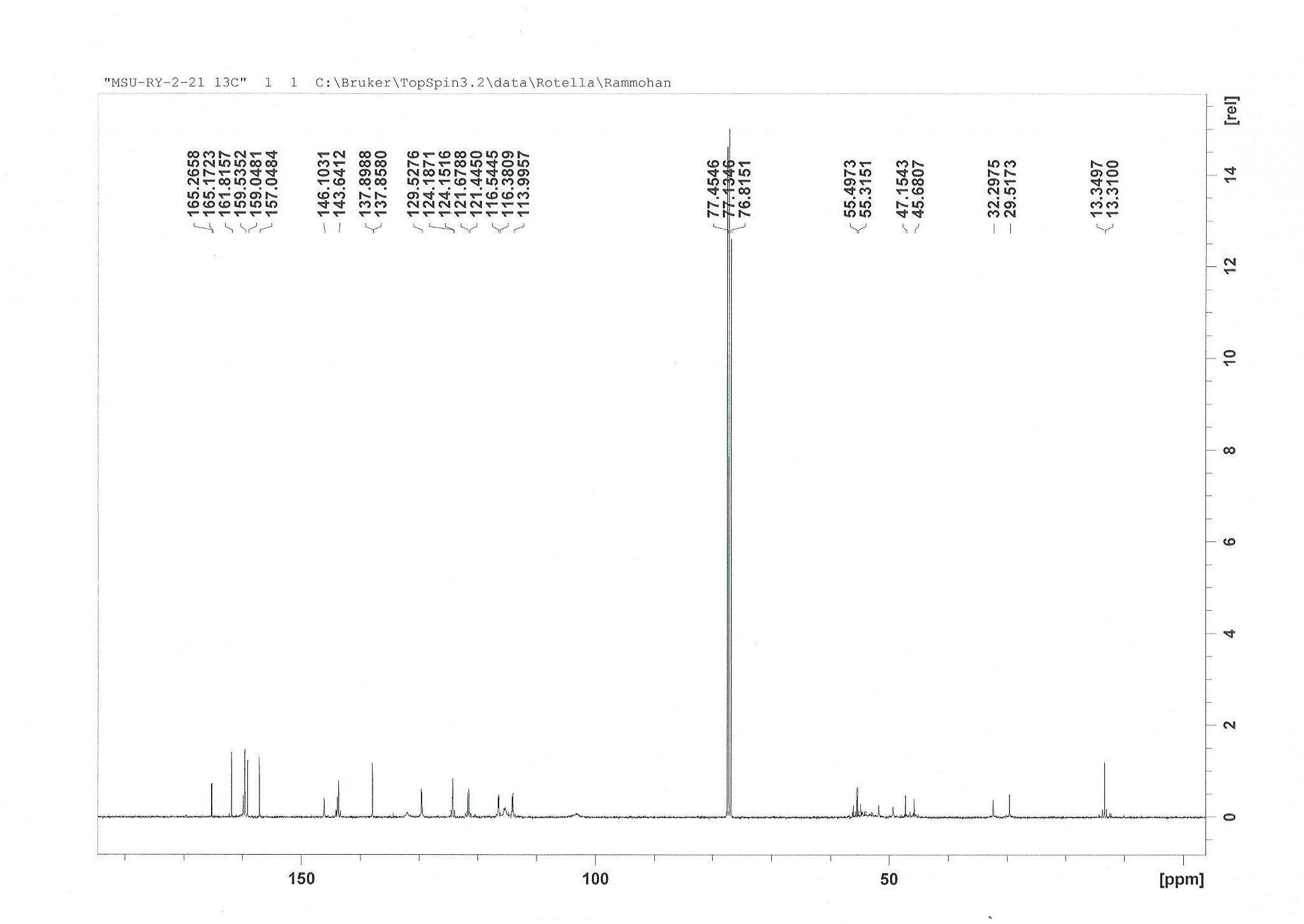

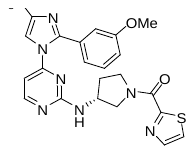

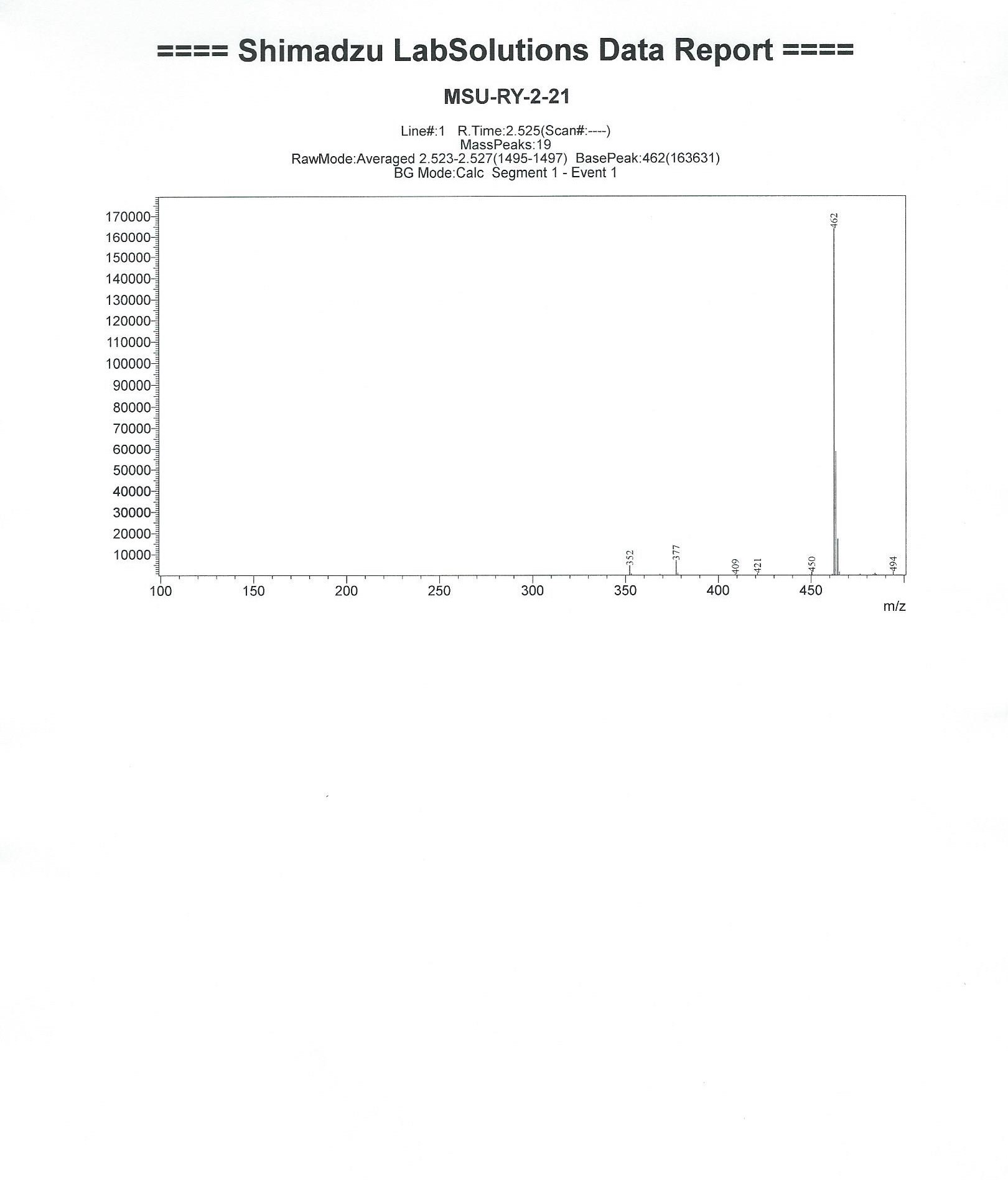

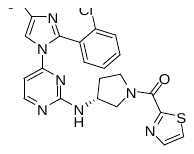

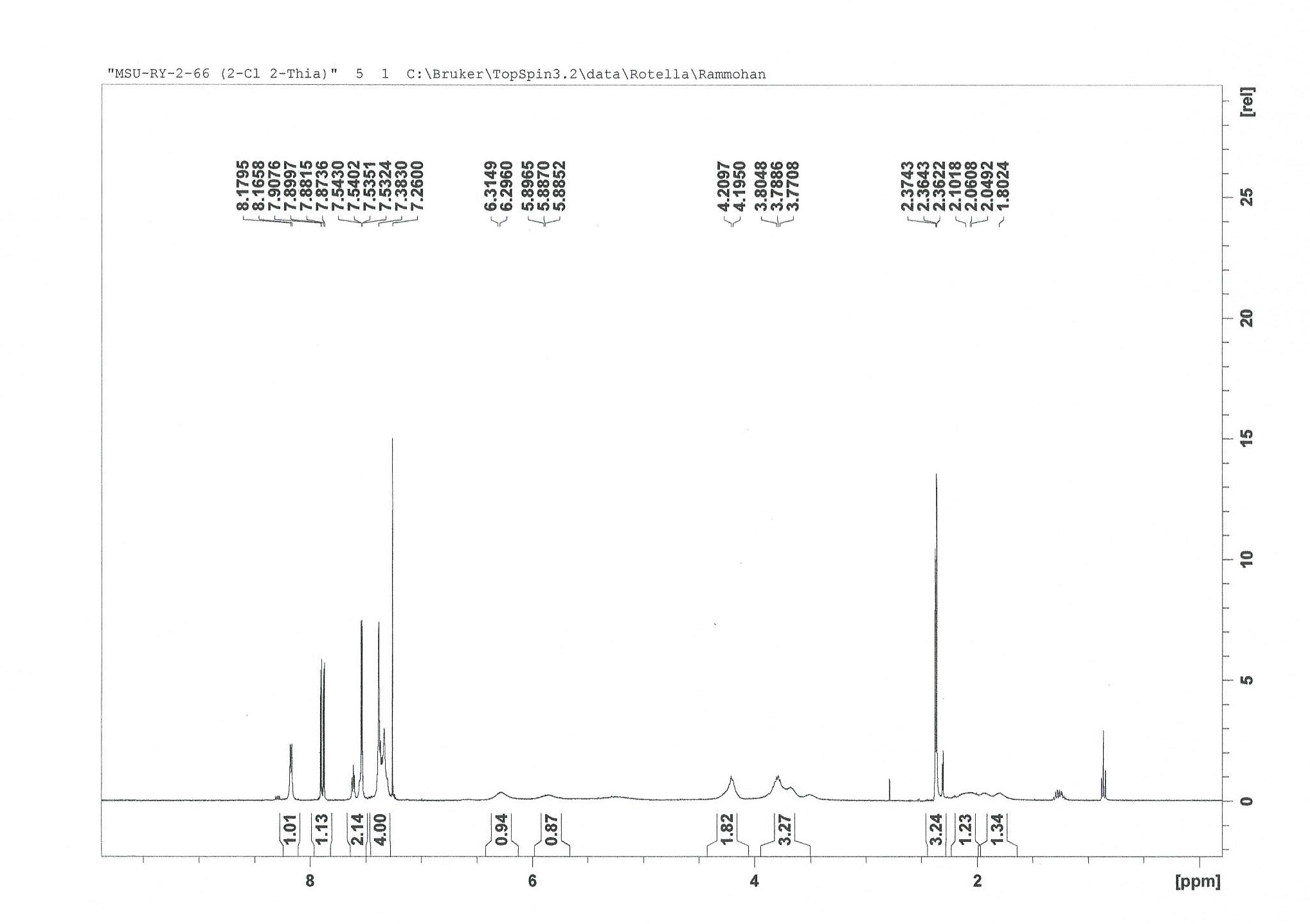

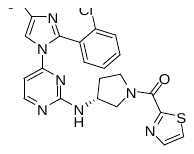

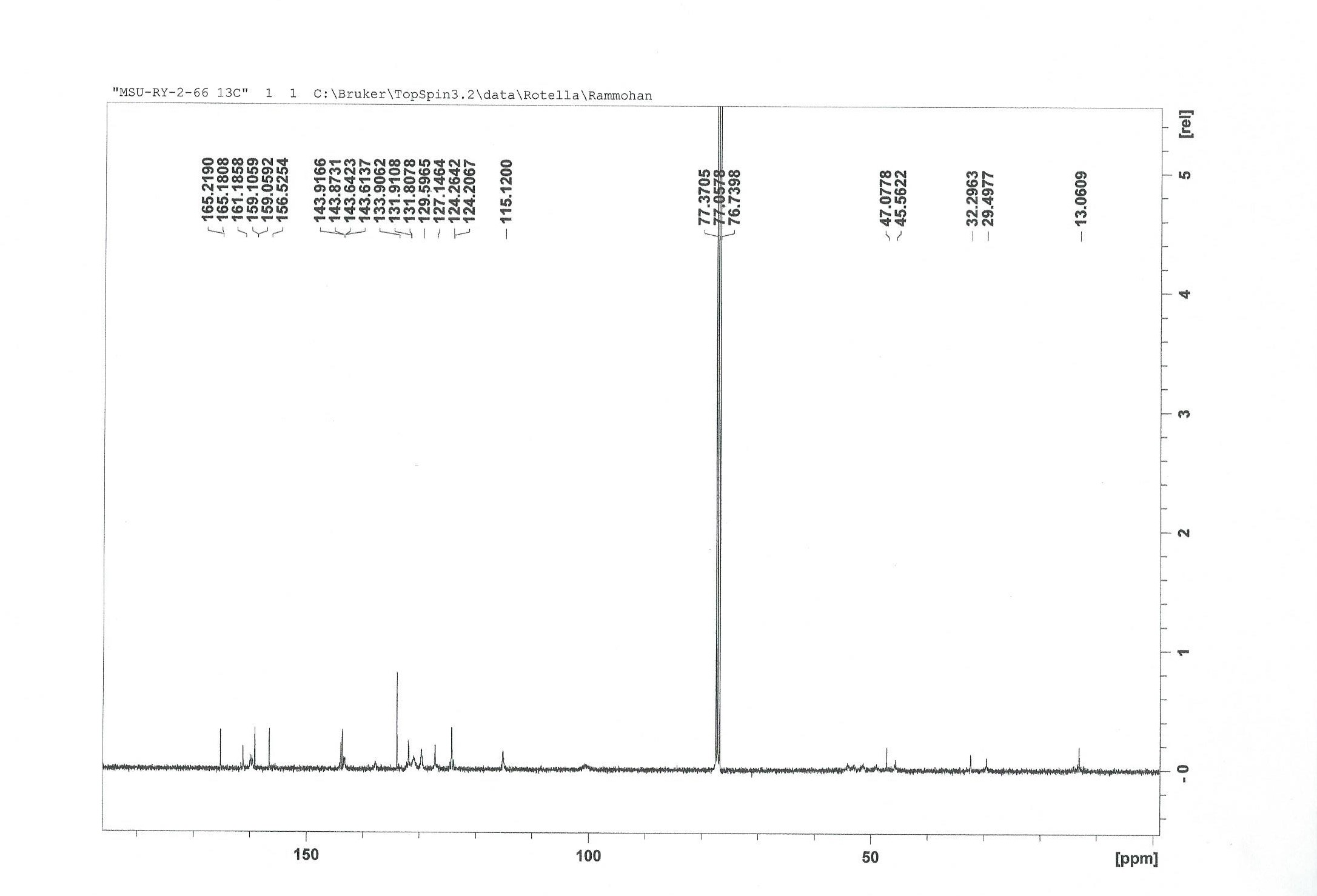

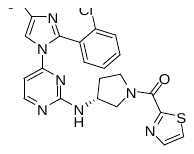

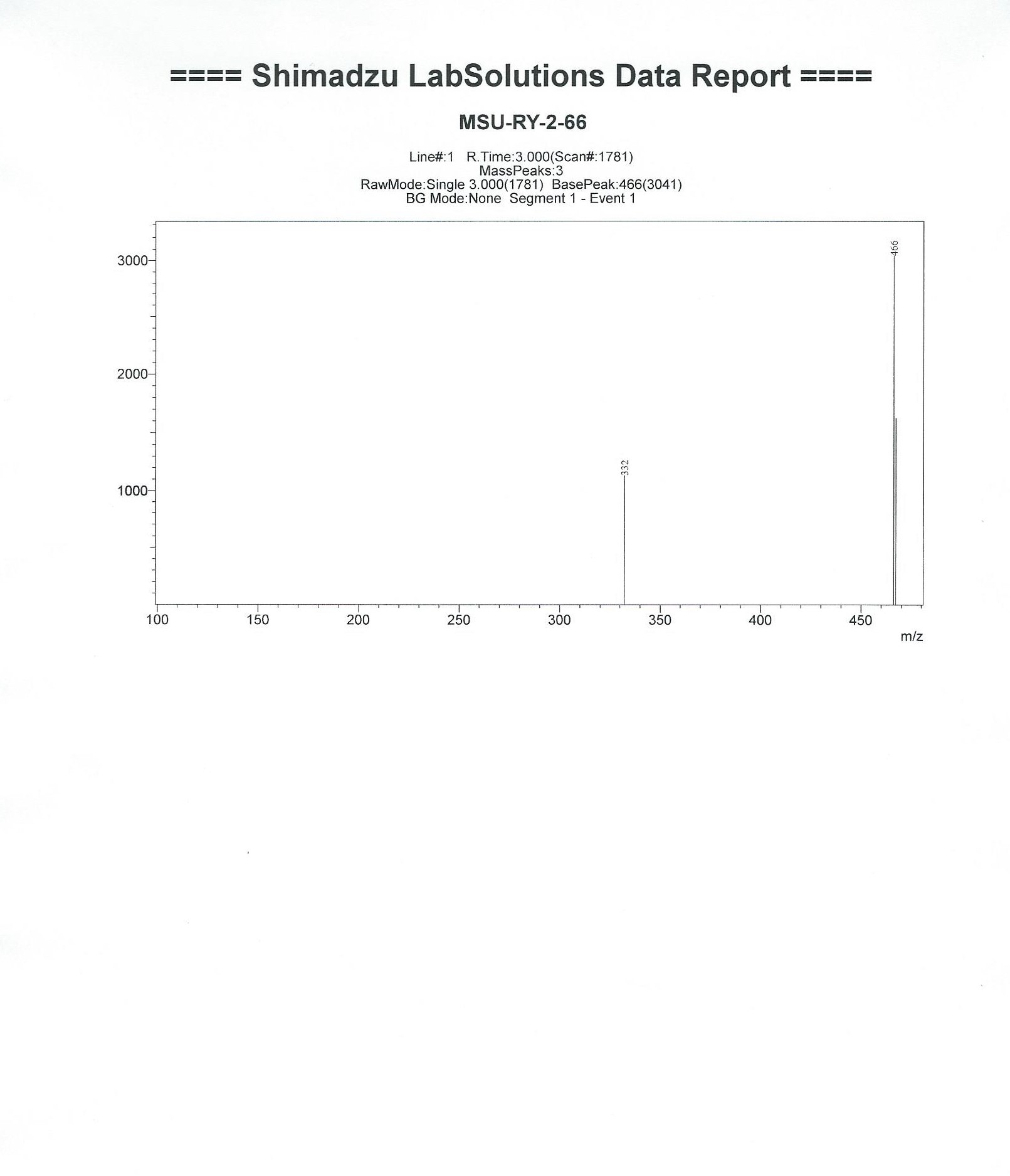

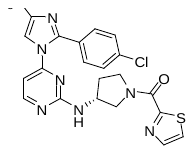

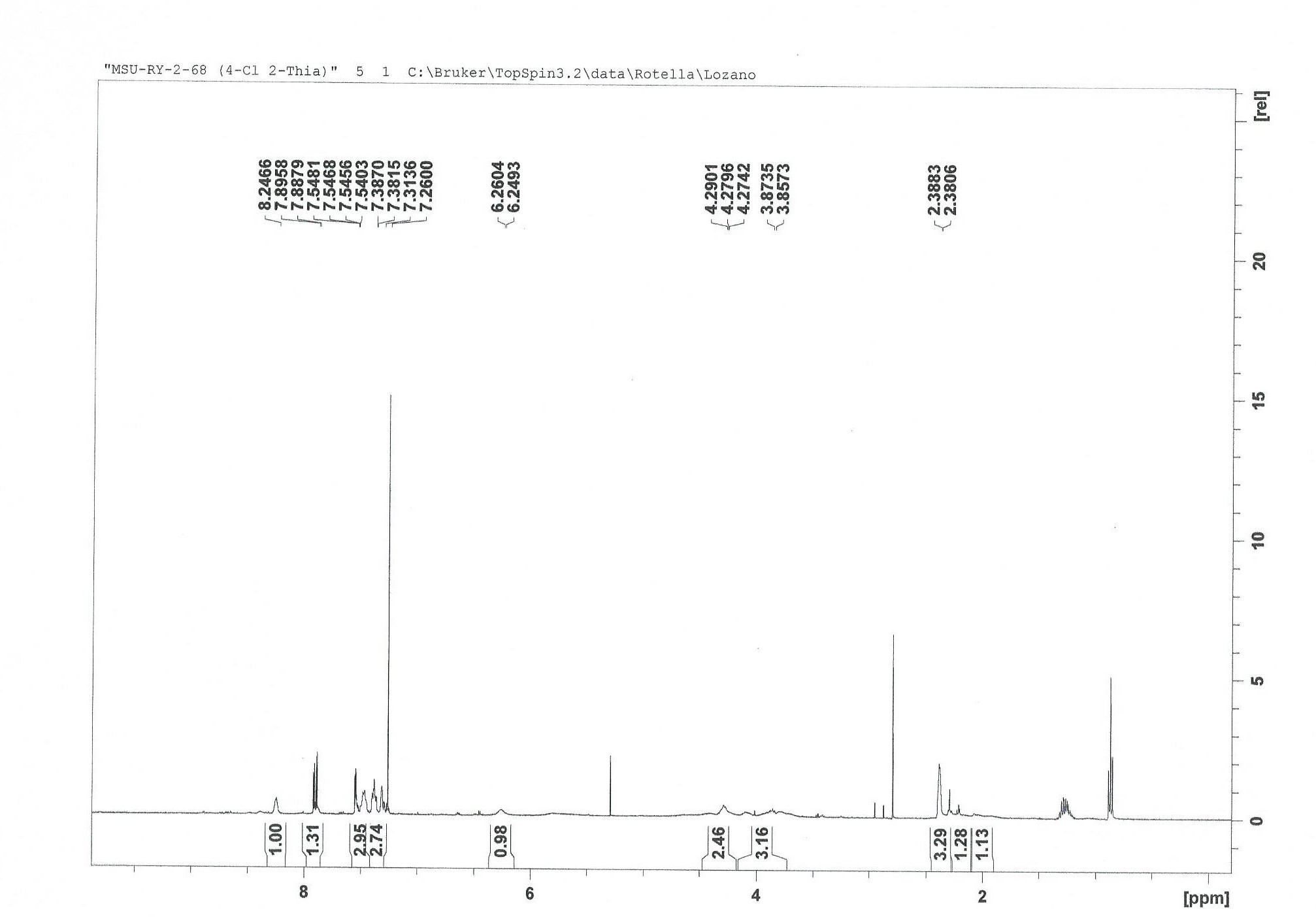

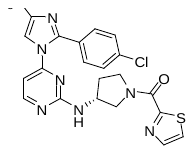

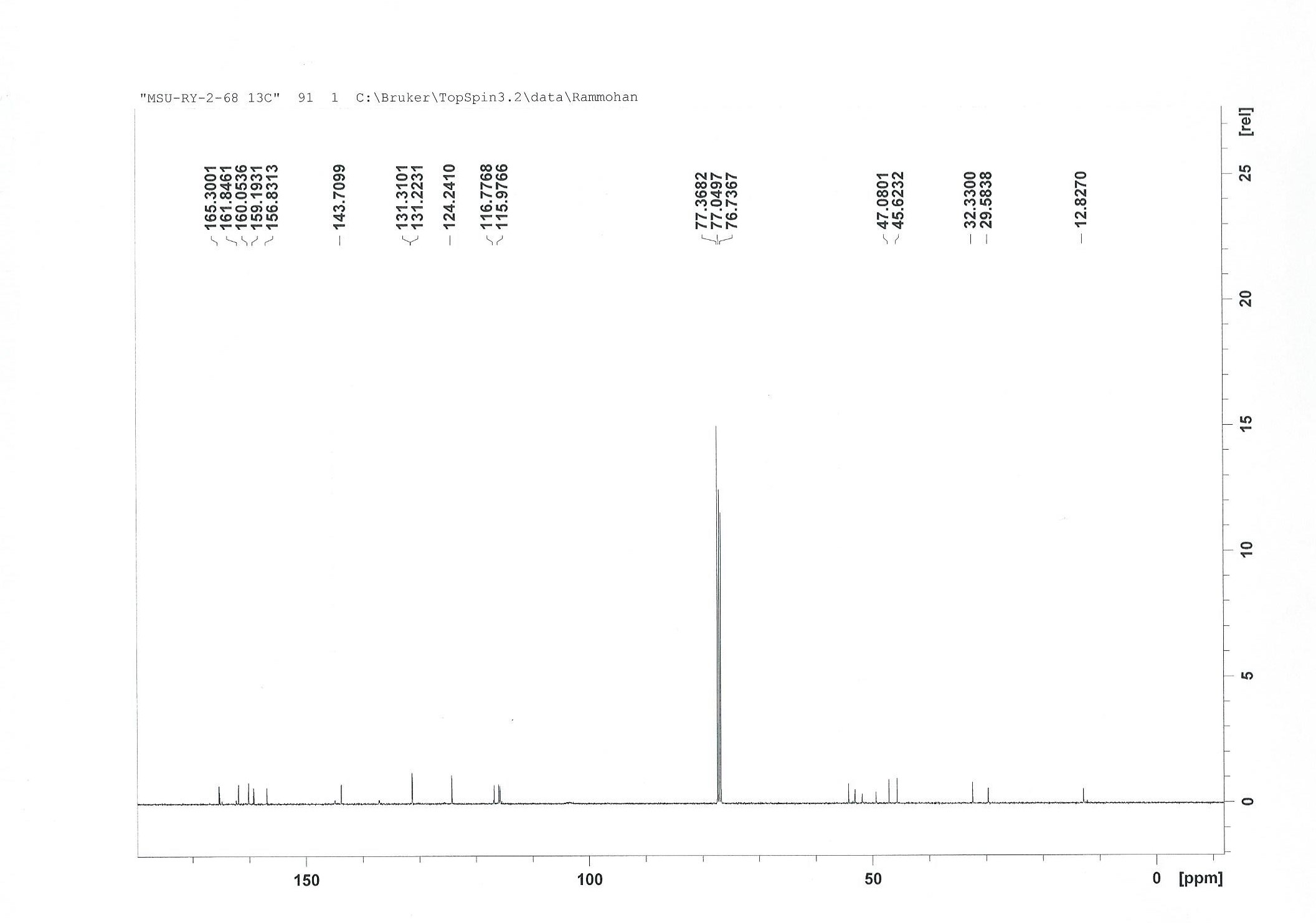

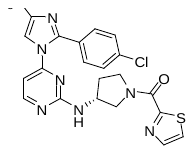

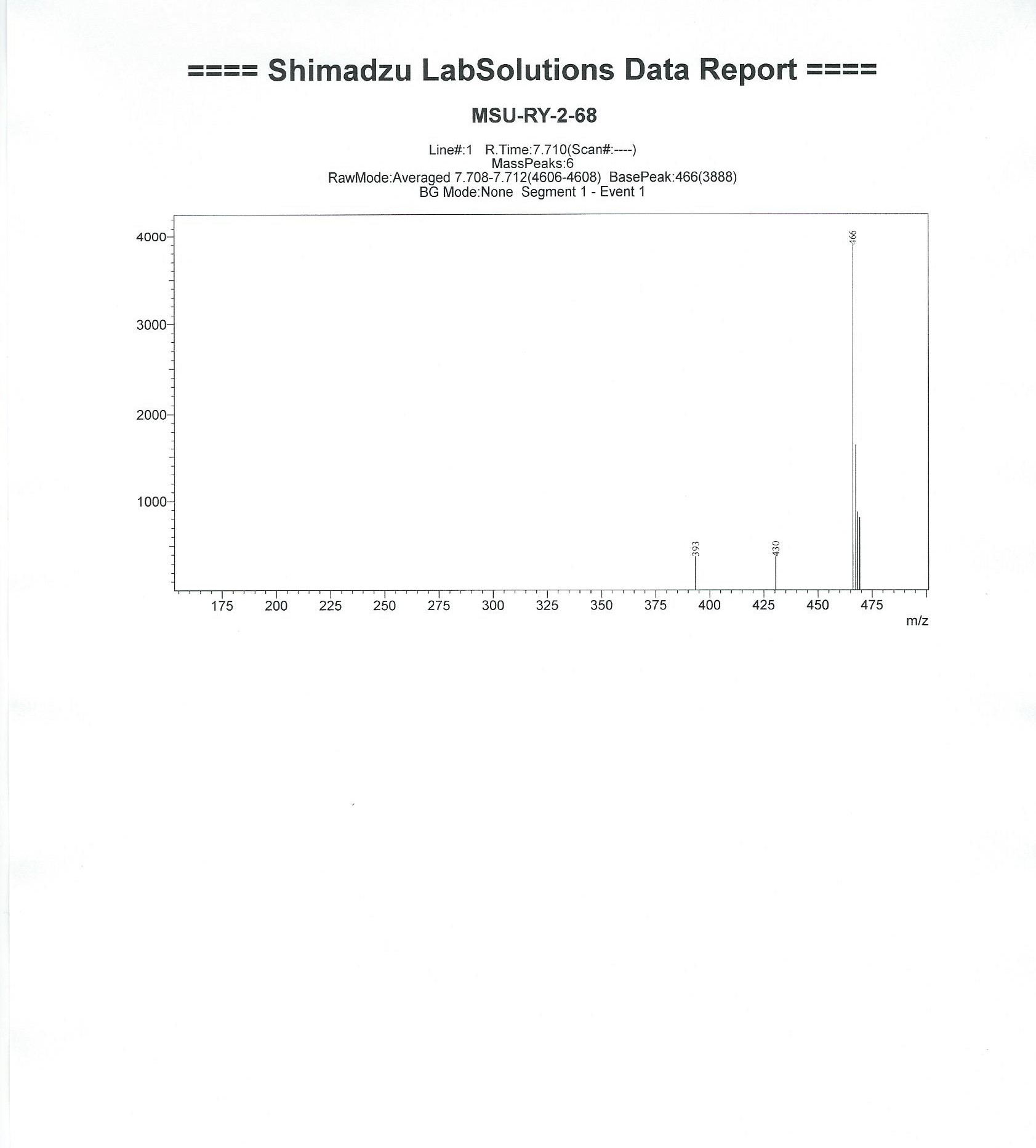

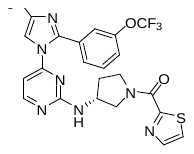

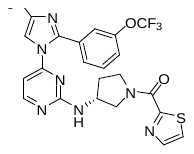

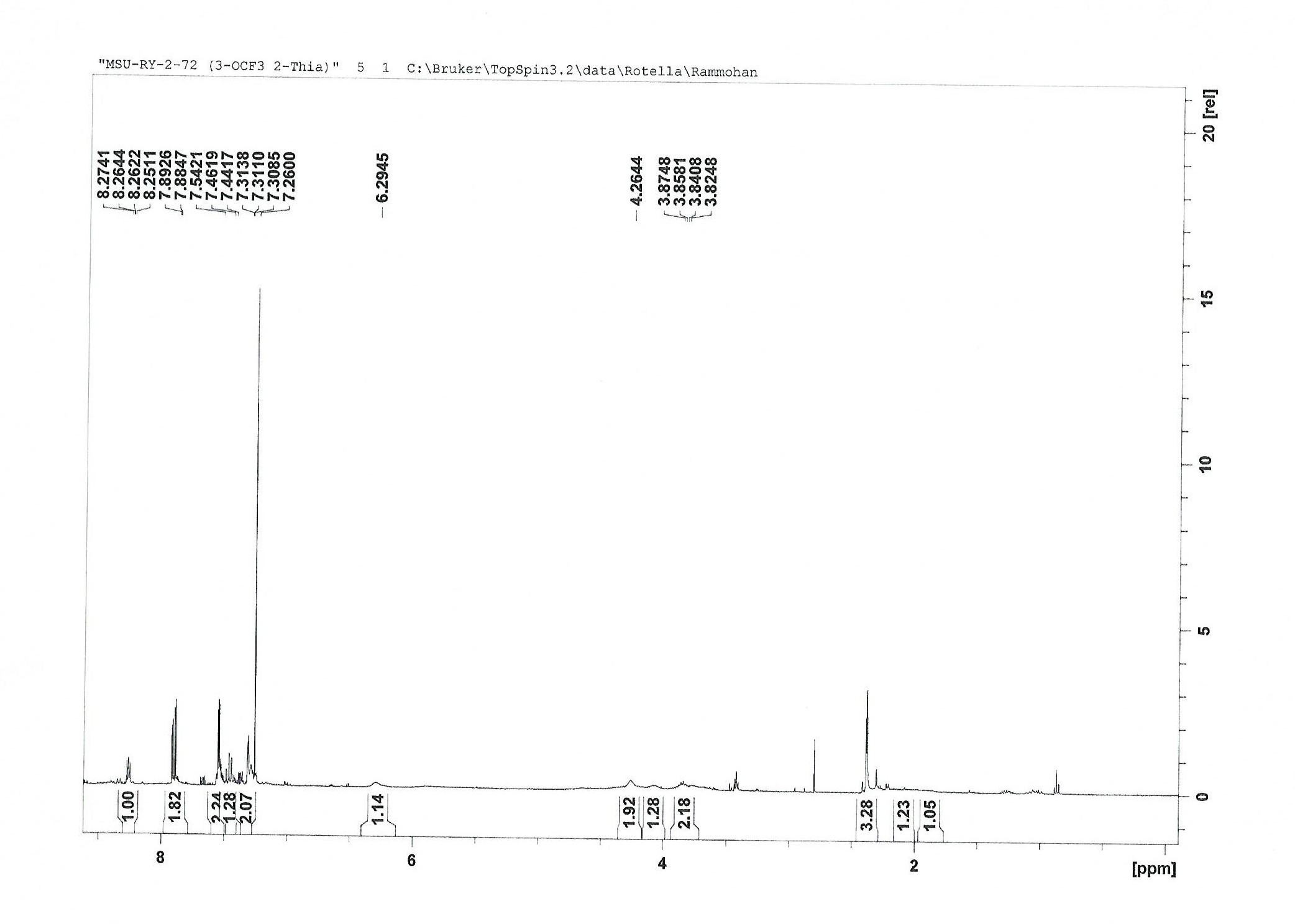

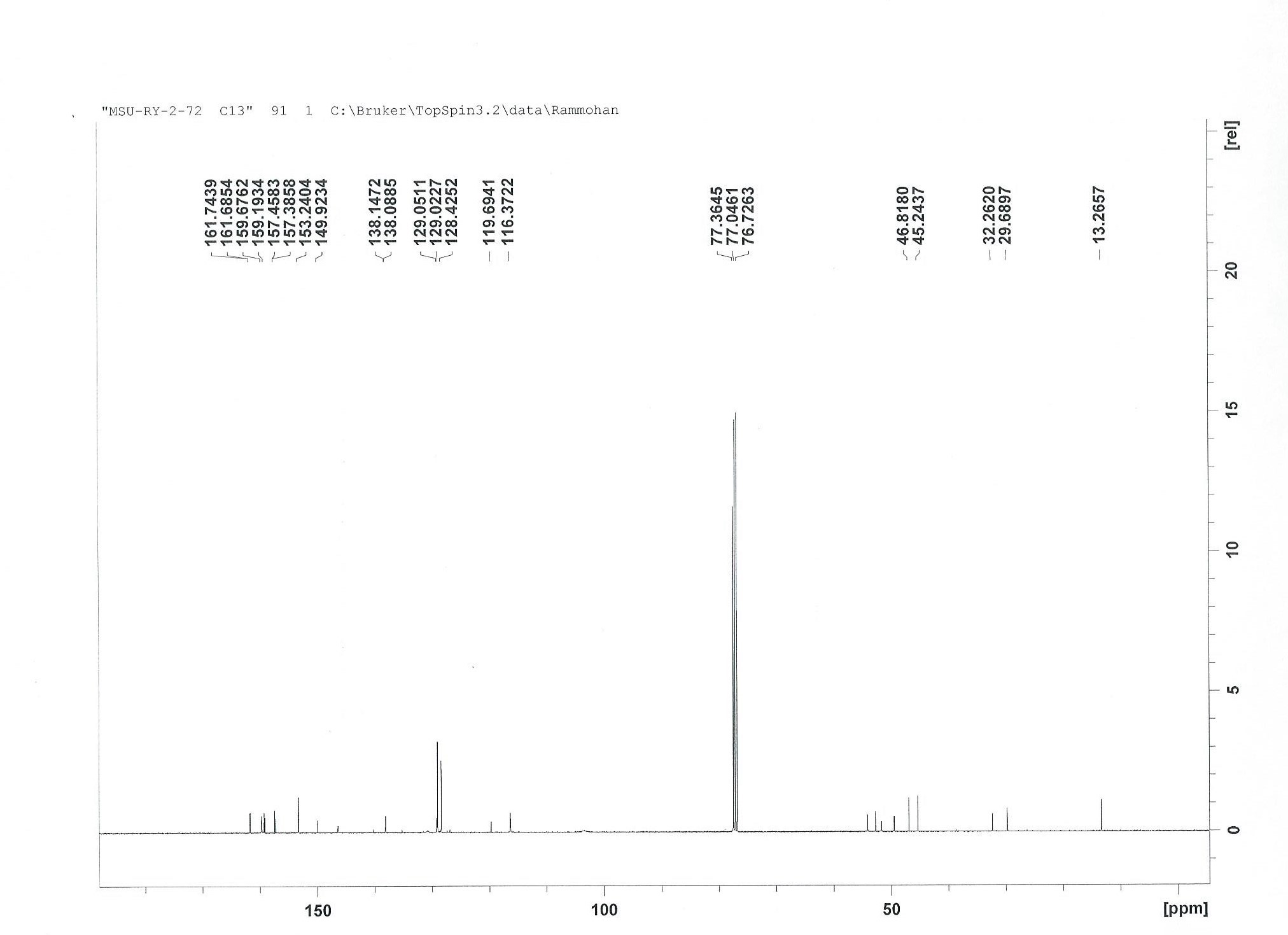

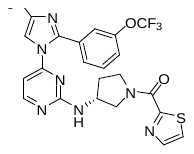

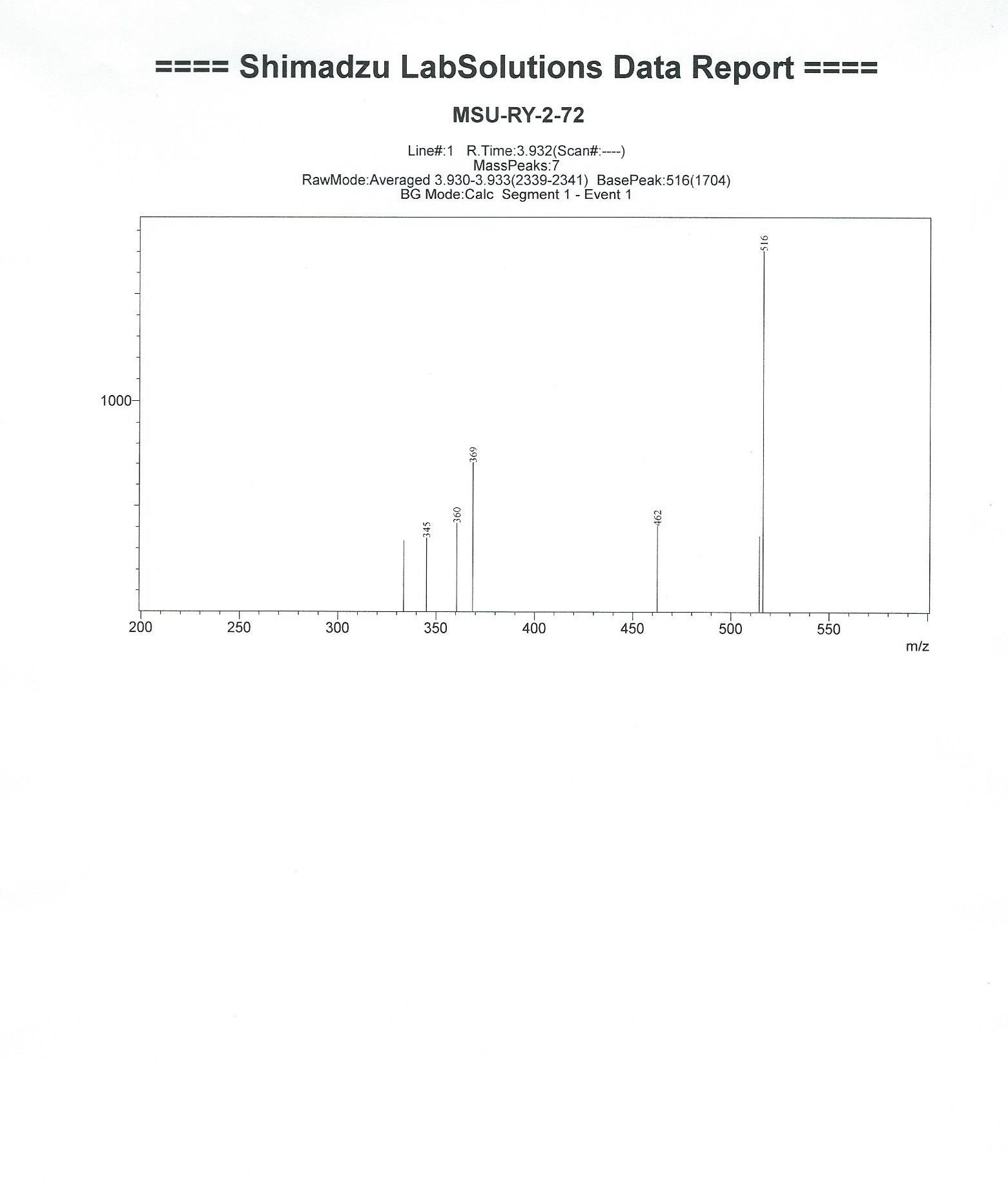
